# HDGF supports anti-apoptosis and pro-fibrosis in pancreatic stellate cells of pancreatic cancer

**DOI:** 10.1101/272542

**Authors:** Yi-Ting Chen, Tso-Wen Wang, Tsung-Hao Chang, Teng-Po Hsu, Jhih-Ying Chi, Yu-Wei Hsiao, Chien-Feng Li, Ju-Ming Wang

**Affiliations:** Medical Research Department, Chi Mei Medical Center, Tainan, Taiwan.; Department of Biotechnology and Bioindustry Sciences, College of Bioscience and Biotechnology, National Cheng Kung University, Tainan, Taiwan.; Institute of Bioinformatics and Biosignal Transduction, College of Bioscience and Biotechnology, National Cheng Kung University, Tainan, Taiwan.; Institute of Basic Medical Science, College of Medicine, National Cheng Kung University, Tainan, Taiwan.; Department of Life Sciences, College of Bioscience and Biotechnology, National Cheng Kung University, Tainan, Taiwan.; Department of Pathology, Chi Mei Medical Center, Tainan, Taiwan.; Interbational Research Center for Wound Repair and Regeneration, National Cheng Kung University, Tainan, Taiwan.; Graduate Institute of Medical Sciences, College of Medicine, Taipei Medical University, Taipei, Taiwan.; Graduate Institute of Medical, College of Medicine, Kaohsiung Medical University, Kaohsiung, Taiwan.

**Keywords:** hepatoma-derived growth factor (HDGF), pancreatic stellate cells (PSCs), anti-apoptosis, pro-fibrosis.

## Abstract

Pancreatic cancer is refractory and characterized by extensively surrounding- and intra-tumor fibrotic reactions that are contributed by activated pancreatic stellate cells (PSCs). Activation of PSCs plays a pivotal role for developing fibrotic reactions to affect themselves or pancreatic cancer cells (PCCs). In the current study, we demonstrated that hepatoma-derived growth factor (HDGF) was secreted from transforming growth factor-β1 (TGF-β1)-treated PSCs. We found that HDGF contributed to anti-apoptosis of PSCs and led to synthesis and depositions of extracellular matrix proteins for stabilizing PSCs/PCCs tumor foci. CCAAT/enhancer binding protein δ (CEBPD) responds to TGF-β1 through a reciprocal loop regulation and further activated hypoxia inducible factor-1α (HIF-1α) contributed to up-regulation of *HDGF* gene. It agrees with the observation that severe stromal growth positively correlated with stromal HDGF and CEBPD in pancreatic cancer specimens. Collectively, the identification of TGF-β1-activated CEBPD/HIF-1α/HDGF axis provides new insights for the novel discoveries of HDGF in anti-apoptosis and pro-fibrosis of PSCs and outgrowth of pancreatic cancer cells.

## INTRODUCTION

Pancreatic cancer shows late detection, aggressive growth, early metastasis, resistance to chemotherapeutic drugs and radiotherapy, and strong fibrotic reactions (Habisch, Zhou et al., 2010). Strong fibrotic reactions are also reported, leading to a fibrotic hypo-vascular barrier that surrounds pancreatic tumors, resulting in poor blood perfusion in transplanted tumors in contrast to the normal pancreas (Olive, Jacobetz et al., 2009). Over the last two decades, gemcitabine has been the cornerstone of chemotherapy for all stages of pancreatic cancer and works by targeting during DNA synthesis (Binenbaum, Na’ara et al., 2015). Compared to patients who receive the traditional chemotherapy drug fluorouracil, the median survival duration is only extended from 4.41 to 5.65 months in gemcitabine-treated patients (Burris, Moore et al., 1997). Though an advanced therapy suggest that gemcitabine-based combination therapies improve overall survival from 8.5 to 11.1 months (Conroy, Desseigne et al., 2011, Von Hoff, Ervin et al., 2013), pancreatic cancer is still refractory to chemotherapeutic drugs due to limited drug delivery resulting from extensively surrounding- and intra-tumor fibrotic reactions (Jacobetz, Chan et al., 2013, McCarroll, Naim et al., 2014, Michl & Gress, 2012, Olive et al., 2009).

Pancreatic stellate cells (PSCs) are a subset of pancreatic cancer-associated fibroblasts that regulate the synthesis and degradation of extracellular matrix (ECM) proteins and provide pro-survival signals to tumors (Apte, Pirola et al., 2012, Fu, Liu et al., 2018, Habisch et al., 2010, Masamune, Watanabe et al., 2009). Though TGF-β1 can contribute to cell proliferation and epithelial-to-mesenchymal transition of pancreatic cancer cells, its details including regulation and biological effects in cancer-associated stromal cells remain largely unclear. In response to TGF-β1, activation of PSCs results in enhanced proliferative rates, differentiated transformation into myo-fibroblast-like cells, and increased secretion of ECM proteins, particularly collagens (Apte et al., 2012). As described above, activated PSCs, which are responsible for producing ECM components to develop a fibrotic microenvironment encasing pancreatic cancer tissues, interact with cancer cells and affect cancer cell functions, such as tumor progression, metastasis, and drug resistance (Habisch et al., 2010). Though the hypoxia-induced pro-fibrotic and pro-angiogenic responses in PSCs have been established in pancreatic cancer progression (Erkan, Reiser-Erkan et al., 2009, Masamune, Kikuta et al., 2008), the interaction of PSCs and PCCs in early stage with normoxia remains largely uninvestigated.

However, recent studies have reported a surprising role of fibrosis in pancreatic cancer to restrain tumor growth through depletion or reduction of the stromal fibrotic content using the α-smooth muscle actin-positive (α-SMA) myo-fibroblast-depleted model (Ozdemir, Pentcheva-Hoang et al., 2014) or sonic hedgehog-deficient mouse model (Rhim, Oberstein et al., 2014). These findings suggest that the PSCs-generating fibrotic microenvironment is important not only for passive scaffold architecture but also as an active regulator of tumors. Hence, targeting the interaction between the tumor and stromal tissues has been suggested as a potential strategy for pancreatic tumor treatment (Kota, Hancock et al., 2017, Li, Ma et al., 2012). Increasing evidence suggests that certain secreted soluble proteins promote tumor progression while the status of PSCs switches from quiescent to active.

Transcriptional activation has been implicated in many cell features, including fibrosis. Following tumorigenesis, the cancer cell-faced environment initiates from normoxic conditions. Hypoxia inducible factor-1α (HIF-1α) is an inducible transcription factor that is generally stable under hypoxic conditions (Majmundar, Wong et al., 2010). Although the involvement and importance of HIF-1α in cancer progression in hypoxia has been fully supported, the regulation and participation of HIF-1α under normoxic conditions remain largely uninvestigated. Additionally, transcription factor CCAAT/enhancer binding protein δ (CEBPD) is induced in response to many external stimuli (Ko, Chang et al., 2015). In tumorigenesis, inactivation of CEBPD in certain cancer cells has been suggested to benefit cancer progression (Chuang, Wang et al., 2014, Ko et al., 2015, Ko, Hsu et al., 2008, Li, Tsai et al., 2015). Interestingly, activation of CEBPD in the tumor microenvironment has been suggested to serve a pro-tumor role (Chi, Hsiao et al., 2015, Hsiao, Li et al., 2013). However, the association of CEBPD expression and regulation in stromal fibroblasts, particularly in stellate cells, remains less investigated.

Hepatoma-derived growth factor (HDGF) is detectable in the conditioned medium of the human hepatoma Huh7 cell line (Nakamura, Izumoto et al., 1994) and has been suggested to play a pro-tumor role in the promotion of hepatoma cell proliferation (Enomoto, Nakamura et al., 2015, Kishima, Yamamoto et al., 2002), metastasis (Yamamoto, Tomita et al., 2006), invasion, epithelial-mesenchymal transition (Chen, Kung et al., 2012), and angiogenesis (Shih, Tien et al., 2012). The divergent HDGF functions participate in a broad range of cancer cell activities. In parallel, HDGF is correlated with hepatic fibrogenesis to serve a pro-fibrogenic role by activating the TGF-β1 pathway in hepatocytes of mice that received a bile duct ligation or carbon tetrachloride treatment (Kao, Chen et al., 2010). HDGF protein is also detectable and increased in the secretome of activated PSCs compared to quiescent PSCs (Wehr, Furth et al., 2011). However, to our knowledge, the involvement and effect of HDGF in pancreatic cancer-associated fibrosis, including its role in PSCs, and its associated contribution in cancer progression remains an open question.

In the present study, we demonstrated that HDGF could be secreted from TGF-β1-treated PSCs and contributed to anti-apoptosis and the generation of the ECM protein-rich microenvironment. We further revealed that the consequent activation of transcription factors CEBPD and HIF-1α was involved in transcriptional activation of *HDGF* gene in PSCs. Stromal HDGF and CEBPD positively associated with the severe stromal growth of pancreatic cancers. In addition to providing new insights into HDGF and CEBPD biology in pancreatic cancer, these results also suggested that HDGF and CEBPD could serve as fibrotic markers as well as potent therapeutic targets for developing an inhibitor of fibrotic reactions.

## RESULTS

### HDGF is responsive to TGF-β1 in PSCs, plays anti-apoptotic and pro-fibrotic effect roles in pancreatic cancer

TGF-β1 activates ECM proteins, leading to pancreatic fibrosis through activation of PSCs. TGF-β1 activate PSCs to highly express α-SMA and vimentin and rearrange F-actin, which represent markers for mesenchymal cells (Bachem, Schneider et al., 1998, Menke & Adler, 2002, Vogelmann, Ruf et al., 2001). TGF-β1, known to mediate fibrosis, is also up-regulated and secreted in pancreatic tumors (Sakurai, Sawada et al., 1997, Shields, Dangi-Garimella et al., 2012). *α-SMA* transcripts is reported to respond to TGF-β1 in RLT-PSCs, immortal human PSCs (Jesnowski, Furst et al., 2005). We next verified these fibrogenic markers in TGF-β1-treated RLT-PSCs using reverse transcription polymerase chain reaction (RT-PCR), western blot and immunofluorescence staining. The results showed that TGF-β1 treatment induced expression of α-SMA and vimentin (**Figure 1A**). Moreover, the experimental RLT-PSCs showed a morphological change into a myo-fibroblast-like shape that exhibited F-actin rearrangements (**Figure 1B**). We further verified the expression of HDGF in response to TGF-β1 in PSCs. We observed that the proliferation of RLT-PSCs had no effect at 0.5 ng/mL of TGF-β1 (**Supplementary Figure 1; (Vonlaufen, Phillips et al., 2010)**), but indeed increased the secretion of HDGF in the conditioned medium of TGF-β1-treated RLT-PSCs (**Figure 1C**).

**Figure 1.**
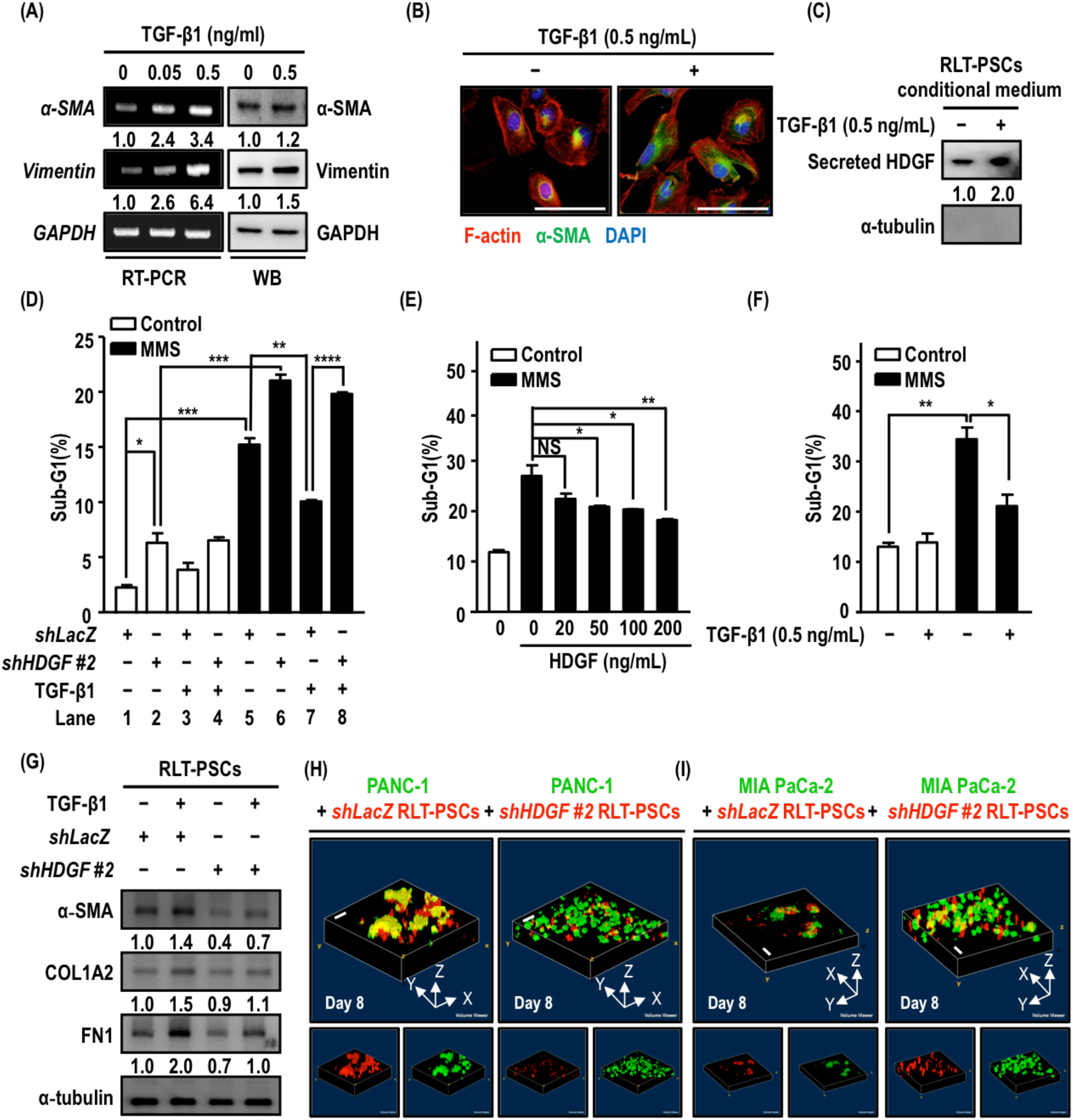
Hepatoma-derived growth factor (HDGF) exerts anti-apoptotic and pro-fibrotic effects of pancreatic stellate cells (PSCs) that interact with pancreatic cancer cells (PCCs). (A) RLT-PSCs, immortal human PSCs, were treated with transforming growth factor-β1 (TGF-β1) in a dose-dependent manner for 24 hours. The fibrogenic markers in TGF-β1-treated RLT-PSCs, α-smooth muscle actin (α-SMA) and vimentin, were further determined using reverse transcription-polymerase chain reaction (RT-PCR) and western blot (WB) assays. (B) Immunofluorescence staining was used to examine the fibrogenic markers in TGF-β1-treated RLT-PSCs, such as the high expression of α-smooth muscle actin (α-SMA) and filamentous actin (F-actin) rearrangement. α-SMA expression and F-actin rearrangement were determined in RLT-PSCs treated with or without TGF-β1 (0.5 ng/mL) for 24 hours. Immunofluorescence staining images were captured under a vertical fluorescent microscope to observe morphological changes (white scale bar 100 μm). Green and red fluorescence indicate expression of α-SMA and F-actin, respectively. (C) The conditioned medium was collected and concentrated from RLT-PSCs with or without TGF-β1 treatment (0.5 ng/mL) for 24 hours. Secreted HDGF expression was determined using western blotting assays, and α-tubulin served as a negative control to exclude the signals from cell lysate of RLT-PSCs. (D) to (F) Methyl methanesulfonate (MMS) treatment was used to induce cell death. Treated RLT-PSCs were harvested and subjected to an analysis of the cell cycle distribution by flow cytometry using propidium iodide staining. The sub-G1 phase indicated the apoptotic cell population. *shLacZ* (control) or *shHDGF #2* (HDGF knockdown) RLT-PSCs were established by lenti-virus delivered short hairpin RNA. *shLacZ*, *shHDGF #2* (D) or parental (E and F) RLT-PSCs were pre-treated with recombined human HDGF as indicated (0, 20, 50, 100, and 200 ng/mL) for 6 hours or TGF-β1 (0.5 ng/mL) for 24 hours. MMS (75 μg/mL) was administered to pre-treated cells for subsequent 24 hours. (G) Expression of α-SMA, type-I collagen (COL1A2), and fibronectin (FN1) was determined in the *shLacZ* and *shHDGF #2* RLT-PSCs under TGF-β1 stimulation (0.5 ng/mL) for 24 hours by western blot assays. (H) and (I) *shLacZ* or *shHDGF #2* RLT-PSCs were respectively transfected with the *mCherry* fluorescent gene (mCherry/*shLacZ* RLT-PSCs or mCherry/*shHDGF #2* RLT-PSCs). PCCs, PANC-1 and MIA PaCa-2 cells, were transfected with the enhanced green fluorescent protein (EGFP) gene (EGFP/PANC-1 or EGFP/MIA PaCa-2 cells). The mCherry/*shLacZ* RLT-PSCs or mCherry/*shHDGF #2* RLT-PSCs were individually co-cultured with EGFP/PANC-1 or EGFP/MIA PaCa-2 cells. The mixed mCherry/RLT-PSCs and EGFP/PCCs (ratio = 5:1) were suspended in 50% matrigel and subjected to a three-dimension system by hanging drop cell culture. After 8 days, the sphere structure was observed at 100× magnification and captured at by confocal microscopy (white scale bar 200 μm). According to the z-section of captured images, the fluorescence intensity was remodeled to reconstruct the three-dimension structure by ImageJ software. The above data are presented as the means ± standard error of the mean from three independent experiments, and the statistical significance was determined using Student’s unpaired T test (^*^ *P* < 0.05; ^**^ *P* < 0.01; ^***^ *P* < 0 .001; ^****^ *P* < 0 .0001).

HDGF influences cancer cells in multifarious functions including transformation, survival and metastasis of cancer cells, and angiogenesis (Bao, Wang et al., 2014). However, the regulation and potential roles of HDGF in PSCs and its consequent effects in pancreatic cancer progression remain unknown. Physically, the increased cell number results from cell proliferation and survival. The involvement of HDGF in the proliferation of fibroblasts was previously suggested (Ooi, Mukhopadhyay et al., 2010); our current results further demonstrated that attenuation of HDGF could induce apoptosis of RLT-PSCs (**Figure 1D, lanes 1 and 2;** the knockdown efficiency of HDGF in *shHDGF* RLT-PSCs was shown in **Supplementary Figure 2A**). We next examined whether HDGF contributed to the anti-apoptosis in PSCs. Methyl methanesulfonate (MMS), a DNA damage inducer and activator of apoptosis, was applied to induce cell apoptosis in RLT-PSCs. We found that HDGF-treated RLT-PSCs were resistant to MMS-induced cell apoptosis (**Figure 1E**). Following the confirmation that TGF-β1 could suppress MMS-induced apoptosis of RLT-PSCs (**Figure 1F**), we found that the anti-apoptotic effect was attenuated in *shHDGF* RLT-PSCs (**Figure 1D, lanes 5 to 8; Supplementary Figure 3A**). These results suggested that HDGF is required for TGF-β1-induced anti-apoptotic effect in MMS-treated RLT-PSCs. Additionally, inhibition of HDGF also attenuated expression of TGF-β1-induced fibrogenic markers, α-SMA, type-I collagen (COL1A2) and fibronectin (FN1) (**Figure 1G; Supplementary Figure 3B**). The results suggested that HDGF, at least in part, contributes to anti-apoptosis and ECM protein synthesis in PSCs.

Following pancreatic cancer progression, fibrosis increases in parallel with the growth of pancreatic cancer. The results described above suggested that HDGF contributes to survival and ECM protein synthesis of PSCs. PSCs have been suggested to accompany and closely interact with PCCs (Suetsugu, Snyder et al., 2015). We next assessed the effect of HDGF in the cell-cell interaction of PSCs and PCCs. To address this issue, RLT-PSCs and PCCs were co-cultured using a hanging drop cell culture system. mCherry-bearing *shLacZ* RLT-PSCs or *shHDGF* RLT-PSCs (mCherry/*shLacZ* RLT-PSCs or mCherry/*shHDGF* RLT-PSCs; **Supplementary Figure 2B**) were individually co-cultured with EGFP-bearing PCCs (EGFP/PANC-1 or EGFP/MIA PaCa-2 cells; **Supplementary Figure 2C**). Regarding the z-sections of the captured images, compared with mCherry/*shLacZ* RLT-PSCs, the constituted images showed that the co-localizing signals of the composite spheres were diminished when EGFP/PANC-1 or EGFP/MIA PaCa-2 cells were co-cultivated with mCherry/*shHDGF* RLT-PSCs (**Figure 1H and 1I; Supplementary Figure 3C and 3D; Supplementary Movie 1 and 2**). These observations suggested that HDGF is able to stabilize PSCs/PCCs tumor foci.

### HDGF in PSCs orchestrates the fibrotic reactions and influences the growth of PCCs

The results suggested that HDGF is involved in generating an ECM protein-rich microenvironment resulting in stabilizing the sizes of PSCs/PCCs tumor foci. A previous study has demonstrated that dense fibrotic reactions serve as a hypo-vascular barrier (Olive et al., 2009). Therefore, the penetration of endothelial cells of blood vessels is appropriate for assessing tissue density. We performed co-implantations of PSCs and PCCs into NOD-SCID mice to determine the essentiality of HDGF in PSCs-generated fibrotic reactions *in vivo*. Control *shLacZ* or *shHDGF* RLT-PSCs were individually mixed with EGFP/PANC-1 or EGFP/MIA PaCa-2 cells. To evaluate the fibrotic reactions, picrosirius red staining was performed to examine the abundance of collagens in tissue sections from xenografted tumors of co-implantations of RLT-PSCs and PCCs. The results showed that the abundance of collagens was reduced in xenografted tumor sections of co-implanted *shHDGF* RLT-PSCs and EGFP/PANC-1 or EGFP/MIA PaCa-2 cells (**Figure 2A and 2B**). The results suggested that HDGF is not only involved in ECM protein synthesis *in vitro*, but also in ECM protein depositions *in vivo*. Additionally, we performed immunofluorescence assays to determine the distribution of subcutaneously co-implanted RLT-PSCs and EGFP/PANC-1 or EGFP/MIA PaCa-2 cells. Compared with the GFP-negative region of xenografted tumors composed of *shLacZ* RLT-PSCs, the mean fluorescence intensity of the COL1A2 and FN1 signals was attenuated in the GFP-negative region of xenografted tumors composed of *shHDGF* RLT-PSCs (**Figure 2C to 2F**). The COL1A2 and FN1 signals was also assessed in the whole tissue sections, and the results showed that both of COL1A2 and FN1 expression was decreased in the tissue sections composed of *shHDGF* RLT-PSCs and EGFP/PANC-1 or EGFP/MIA PaCa-2 cells (**Figure 2C to 2F**). These observation was consistent with the results from picrosirius red staining tissue sections (**Figure 2A to 2B**). Above results indicated that the diminished depositions of type I collagen and fibronectin associate with the disruption of fibrotic reactions when *shHDGF* RLT-PSCs are co-cultivated with PCCs. Interestingly, expression of CD31, an angiogenesis-related endothelial cell marker, was enhanced in the xenografted tumor sections of co-implanted *shHDGF* RLT-PSCs and EGFP/PANC-1 or EGFP/MIA PaCa-2 cells (**Figure 2G and 2H**), suggesting that the essentiality of HDGF involves in a fibrotic hypo-vascular barrier and results in attenuating the penetration of blood vessels. This result is consistent with the suggestion that HDGF exerts a pro-fibrotic effect to orchestrate the fibrotic reactions. Additionally, *in vivo* imaging system (IVIS) spectrum was applied to evaluate xenografted tumor growth of co-implanted RLT-PSCs and EGFP/PANC-1 or EGFP/MIA PaCa-2 cells expressing green fluorescent protein. Compared to individual xenografted tumor of co-implanted *shLacZ* RLT-PSCs and EGFP-bearing PCCs, co-implantations of *shHDGF* RLT-PSCs and EGFP-bearing PCCs showed larger tumor sizes (**Figure 2I and 2J**). Furthermore, the status of PCCs in the co-implantations of PSCs and PCCs *in vivo* animal study was pathologically evaluated using H&E staining. The higher levels of mitotic activity and more apoptotic bodies were observed in PCCs of co-implanted *shHDGF* RLT-PSCs and EGFP/PCCs *in vivo* (**Supplementary Figure 4**). These observations suggested PCCs own a more rapid mitotic rate and a higher apoptotic index when co-implanted with *shHDGF* RLT-PSCs that results in less fibrotic reactions. In contrast, PCCs co-implanted with *shLacZ* RLT-PSCs show lower resistance to apoptosis and less mitotic activity within a higher fibrotic background. Collectively, our results suggested that *HDGF-*deficient PSCs results in restraints of fibrotic reactions and enhance cancer cell outgrowth.

**Figure 2.**
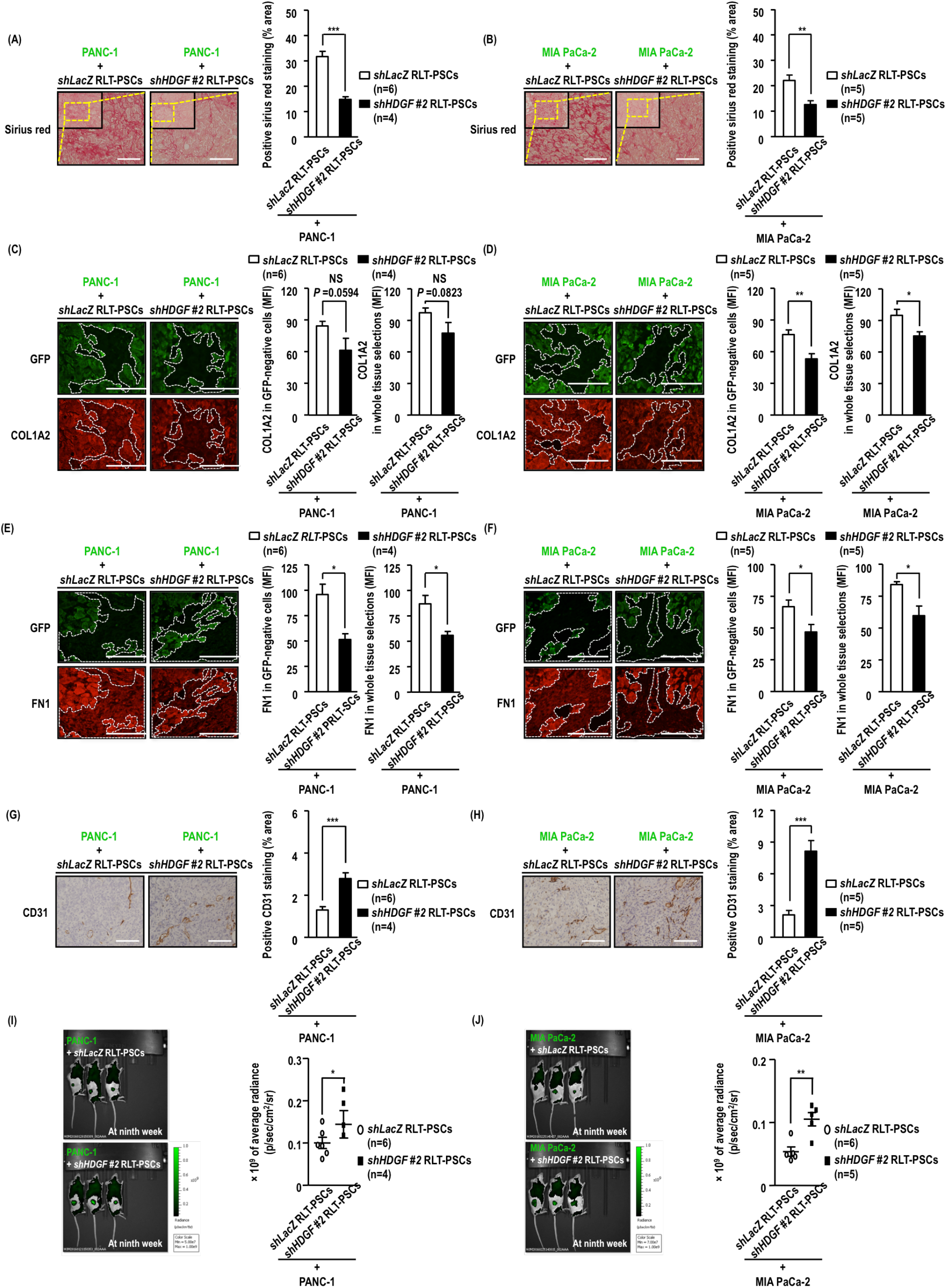
HDGF knockdown PSCs attenuate fibrotic effects and influence the outgrowth of PCCs *in vivo.* *shLacZ* RLT-PSCs or *shHDGF #2* RLT-PSCs were individually mixed with EGFP/PCCs (EGFP/PANC-1 or EGFP/MIA PaCa-2 cells). The mixture of RLT-PSCs and EGFP/PCCs (ratio = 1:1) containing with 50% matrigel was subcutaneously injected into non-obese diabetic-severe combined immunodeficiency mice. Each group contained 4 to 6 mice. After mice were sacrificed at the ninth week, tissue sections of subcutaneous tumors were subjected to immunofluorescence, Picrosirius red staining, and immunohistochemistry analysis. (A) and (B) Picrosirius red staining was applied to evaluate the expression of collagens. The images (left-upper panels) were observed at 100× magnification; the areas bound by the yellow dotted line were observed at 200× magnification under a vertical microscope and the captured images were shown as the right-lower panels (white scale bar 100 μm). Red color indicates positive staining of collagens, which are quantified in the right panels. (C) to (F) Expression of COL1A2, FN1, and green fluorescent protein (GFP) was determined using immunofluorescence stain. The images were observed at 400× magnification under a vertical microscope (white scale bar 100 μm). GFP expression in the upper panels indicates the locations of PCCs. The white dished line circles the regions of GFP-negative cells representing stromal cells. The mean fluorescence intensity (MFI) of COL1A2 and FN1 in GFP-negative cells are quantified by ImageJ software and shown in the right panels. (G) and (H) Immunohistochemistry assays were performed to determine the tumor vasculature by staining with the endothelial cell marker CD31. The images were observed at 200× magnification (white scale bar 100 μm); the brown color indicates positive staining of CD31 expression, which is quantified in the right panels. (I) and (J) IVIS spectrum was applied to evaluate tumor growth in co-implantations of RLT-PSCs and PCCs according to the intensity of green fluorescence expressing in EGFP/PANC-1 or EGFP/MIA PaCa-2 cells. Above results are presented as the means ± standard error of the mean; the statistical significance was determined using Student’s unpaired T test (NS stands for not significant; ^*^ *P* < 0.05; ^**^ *P* < 0.01; ^***^ *P* < 0 .001; ^***^ *P* < 0.0001).

### TGF-β1/CEBPD/ HIF-1α axis contributes to HDGF transcriptional activation in PSCs

We found that TGF-β1-induced HDGF production was detectable in both cell lysates and the conditioned medium from RLT-PSCs by confirming with western blotting and ELISA assay, respectively (**Figure 1C, 3A and 3B**). The results indicated that HDGF production is synchronous with its transcriptional activation. Following the observation that HDGF production is responsive to TGF-β1 in PSCs, using open access websites TFSEARCH (version 1.3) and PROMO (version 3.0.2), we found that *HDGF* promoter contains putative HIF-1α binding motifs. We therefore first examined whether HIF-1α was responsive to TGF-β1 in RLT-PSCs. As well as HDGF expression, both *HIF-1α* mRNA and HIF-1α protein were up-regulated in TGF-β1-treated RLT-PSCs (**Figure 3A**). Though HIF-1α is an inducible transcript factor and tends towards stabilization under a hypoxic condition that have been suggested, we were interested to examine whether TGF-β1 could stabilize HIF-1α under a normoxic condition. We found that TGF-β1 had no effect on HIF-1α protein stability (**Supplementary Figure 5**). Moreover, reporter and chromatin immunoprecipitation (ChIP) assays showed that HIF-1α was responsive to TGF-β1 and could activate the *HDGF* reporter and directly bound to the *HDGF* promoter region under a normoxic condition (**Figure 3C and 3D**). Regarding the idea that the interaction of cancer and stellate cells should be initiated from a normoxic condition, these results led us to test whether TGF-β1 could up-regulate *HIF-1α* transcription in normoxia.

**Figure 3.**
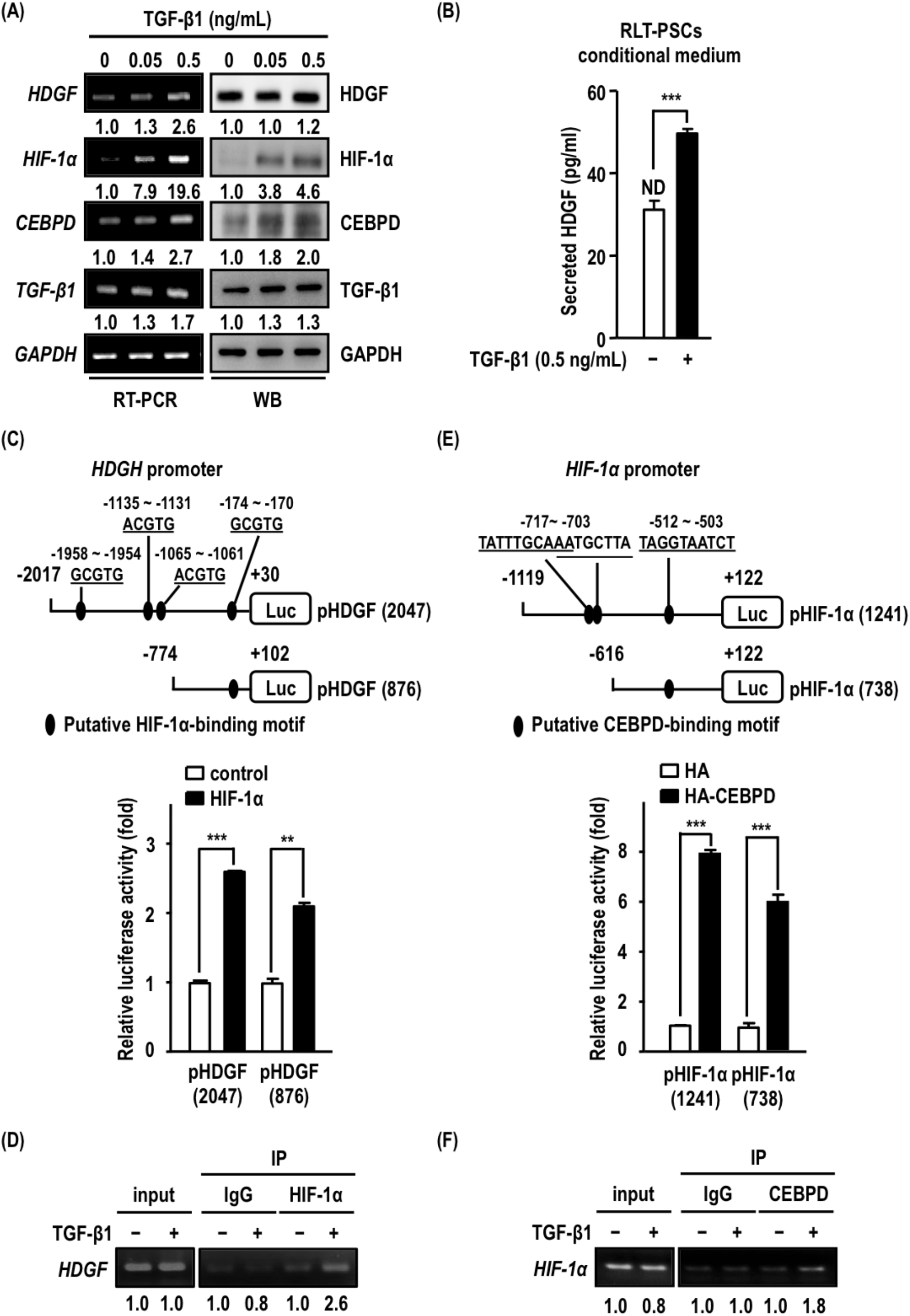
HDGF is transcriptionally up-regulated by TGF-β1/ Hypoxia inducible factor-1α (HIF-1α)/ CCAAT/enhancer binding protein δ (CEBPD) axis. (A) After 0.5 ng/mL of TGF-PS treatment for 24 hours, the mRNA or protein levels of HDGF, HIF-1α, CEBPD, and TGF-β1 were determined using RT-PCR or western blot assays in RLT-PSCs, respectively. (B) After 0.5 ng/mL of TGF-β1 treatment for 24 hours, the levels of HDGF in conditioned medium containing 1% bovine serum albumin, were determined using enzyme-linked immunosorbent assay. (ND is represented as not detectable; the minimum detectable concentration is 30 pg/mL). (C) The putative HIF-1α binding motifs on *HDGF* promoter region were shown in upper panel. The HDGF reporter vector was transfected into RLT-PSCs simultaneously with the pCEP4/HIF-1α expressing vector (or pcDNA3/HA expressing vector as a control group) for 18 hours; then, a luciferase reporter assay was performed to confirm the putative HIF-1α-binding motifs on *HDGF* promoter region. The results are from three independent experiments and presented as the means ± standard error of the mean; the statistical significance was determined using Student’s unpaired T test (^***^ *P* < 0 .001). (D) A chromatin immunoprecipitation assay was performed to determine the binding of HIF-1α onto the *HDGF* promoter region in RLT-PSCs treated with 0.5 ng/mL TGF-β1 for 24 hours. The IgG antibody was used as a negative control. (E) A luciferase reporter assay was used to confirm the putative CEBPD-binding motifs on *HIF-1α* promoter region. The putative CEBPD binding motifs on *HIF-1α* promoter region were shown in upper panel. RLT-PSCs were transfected with the *HIF-1α* reporter vector and pcDNA3/HA or pcDNA3/CEBPD expressing vectors for 18 hours and subsequently subjected to luciferase reporter assays. The results are presented as the means ± standard error of the mean, and the statistical significance was determined using Student’s unpaired T test (^**^ *P* < 0.01; ^***^ *P* < 0 .001). (F) The results of the chromatin immunoprecipitation assays showed the interaction between CEBPD and the *HIF-1α* promoter in RLT-PSCs treated with 0.5 ng/mL TGF-β1 for 24 hours. The above data are from three independent experiments.

Using the same strategy, the putative CEBPD binding motifs was predicted on *HIF-1α* promoter. To examine whether CEBPD responded to TGF-β1 stimulation and regulated *HIF-1α* transcription in PSCs, we first examined whether CEBPD was responsive to TGF-β1 treatment. The results showed that TGF-β1 could induce *CEBPD* and auto-regulated *TGF-β1* transcripts and expression (**Figure 3A**). Moreover, the reporter assay showed that exogenous expression of CEBPD activated the *HIF-1α* reporter in RTL-PSCs (**Figure 3E**). Subsequently, a ChIP assay was performed to verify whether CEBPD could bind to the *HIF-1α* promoter upon TGF-β1 stimulation. The results showed that the CEBPD binding on *HIF-1α* promoter was responsive to TGF-β1 in RLT-PSCs (**Figure 3F**).

Regarding the results that the CEBPD/HIF-1α axis contributes to TGF-β1-induced HDGF expression and HDGF contributes to the suppression of MMS-induced apoptosis in PSCs, we therefore assessed whether both CEBPD and HIF-1α played an anti-apoptotic role in TGF-β1-suppressed apoptosis of MMS-treated RLT-PSCs. In *shCEBPD* or *shHIF-1α* RLT-PSCs (**Supplementary Figure 6A and 6B**), TGF-β1 failed to suppress MMS-induced apoptotic activities (**Supplementary Figure 6C to 6F**).

### CEBPD responds to TGF-β1 through a reciprocal loop regulation

As shown above, CEBPD is responsive to TGF-β1 and contributes to HIF-1α expression that consequently activates HDGF abundance in PSCs. We further verified the upstream role of CEBPD via gain-of- and loss-of-function assays. Following exogenously expressing CEBPD in RLT-PSCs, expression of TGF-β1, α-SMA, CEBPD, HIF-1α, and HDGF was induced (**Figure 4A**). By contrast, expression of these proteins was attenuated in *shCEBPD* RLT-PSCs upon TGF-β1 treatment (**Figure 4B; Supplementary Figure 7A**). Interestingly, in addition to up-regulation by TGF-β1, CEBPD also regulated TGF-β1 expression in RLT-PSCs, suggesting a positive feedback regulation between CEBPD and TGF-β1 in RLT-PSCs (**Figure 4A and 4B**). Moreover, TGF-β1-induced F-actin rearrangement, which represents fibroblast-to-myofibroblast transition, was attenuated in *shCEBPD* RLT-PSCs (**Figure 4C; Supplementary Figure 7B**).

**Figure 4.**
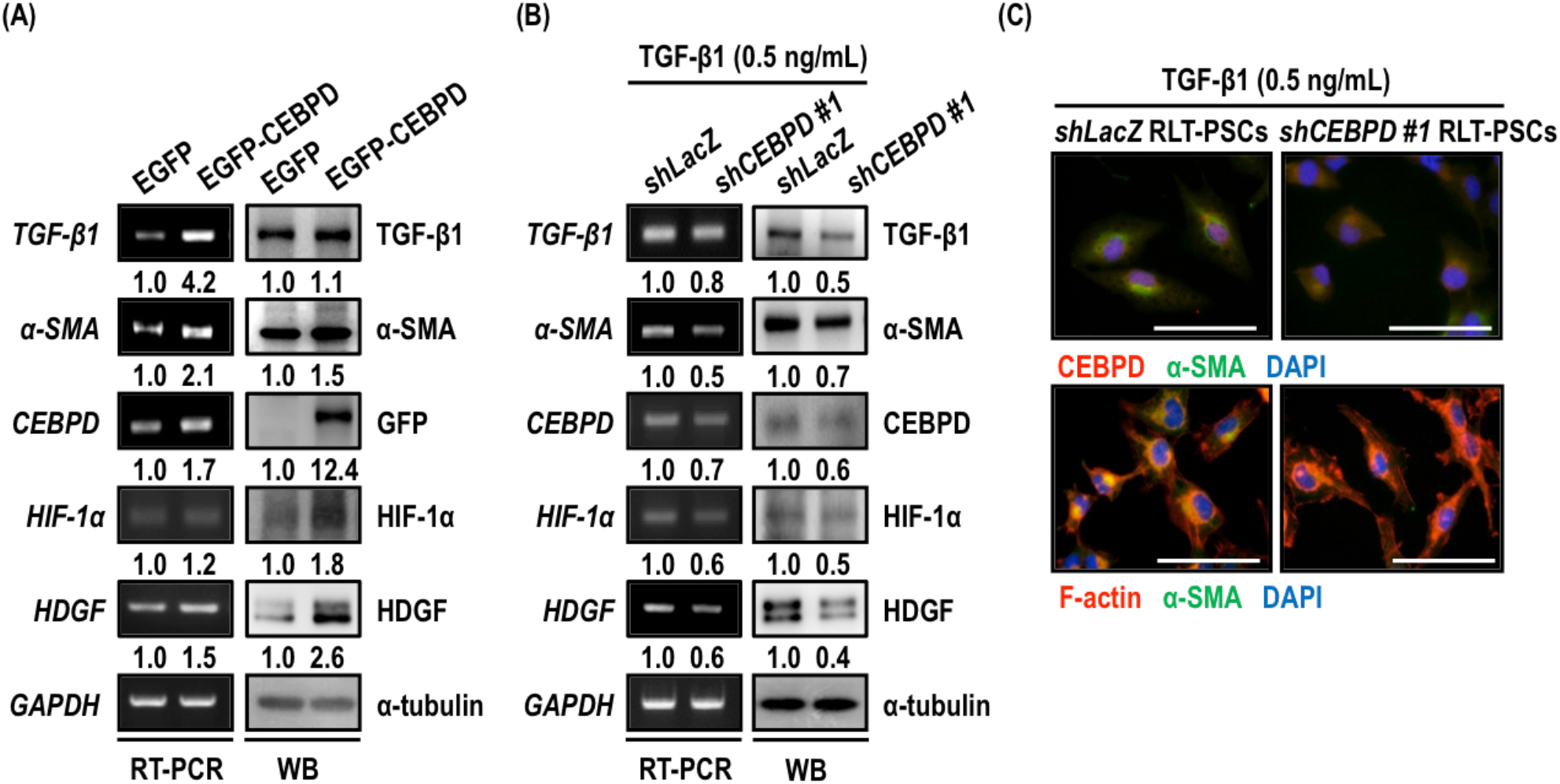
TGF-β1 induces CEBPD expression through a positive feedback loop. (A) After RLT-PSCs were transfected with pcDNA3/EGFP-CEBPD or the pcDNA/EGFP expressing vectors for 18 hours, cells were harvested and subjected to RT-PCR and western blot assays. The mRNA and protein expression levels of α-SMA, CEBPD, HIF-1α, and HDGF were determined in RLT-PSCs with or without CEBPD overexpression. (B) In addition, *shLacZ* RLT-PSCs and *shCEBPD #1* RLT-PSCs were subjected to RT-PCR and western blot assays to examine the mRNA and protein expression levels of α-SMA, CEBPD, HIF-1α, and HDGF after TGF-β1 treatment (0.5 ng/mL) for 24 hours. (C) The fibrogenic markers in TGF-β1-treated RLT-PSCs that could be used to observe morphological changes such as the high expression of α-SMA and F-actin rearrangement, were determined using immunofluorescence staining in *shLacZ* RLT-PSCs or *shCEBPD #1* RLT-PSCs treated with TGF-β1 (0.5 ng/mL) for 24 hours (lower panels). The green and red fluorescence indicates expression of α-SMA and F-actin, respectively. In addition, CEBPD expression was evaluated in treated cells using immunofluorescence staining (upper panels). The green and red fluorescence indicates expression of α-SMA and CEBPD, respectively. The white scale bar indicated 100 μm. The above data are representative from three independent experiments.

### HDGF and CEBPD expression is positively associated with anti-apoptosis and severe stromal growth in human pancreatic cancer specimens

The above results suggested that the TGF-β1-activated CEBPD/HIF-1α/HDGF axis in PSCs contributes to anti-apoptotic and pro-fibrotic effects that severs as a stabilizer of PSCs/PCCs tumor foci. We next assessed the clinical relevance of apoptotic activity, COL1A2, HDGF, and CEBPD expression in stromal cells using a commercial tissue array. According to cell proportion ratio of stromal cells versus total tissue cells, 78 human primary pancreatic cancer specimens were subdivided into mild (<50%) or severe (≥50%) stromal growth. After the expression levels of apoptotic activity, COL1A2, HDGF, and CEBPD in stroma were determined in 78 human pancreatic cancer specimens using immunofluorescence staining, histochemistry-score (H-score) method assigned a score of 0-300 to each specimen for assessing the extent of immunoreactivity by an expert pathologist. The apoptotic activity in stroma negatively associated with the levels of stromal HDGF expression (**Figure 5A brown color and Figure 5B**). Stromal COL1A2 expression showed strong association (*P*<0.001) with severe stromal growth (**Figure 5A red color and 5C**). We also observed significant association between severe stromal growth and high HDGF or CEBPD expression in stroma (**Figure 5A green and cyan colors**, **Figure 5D and 5E**). According to Tumor/Node/Metastasis (TNM) staging of pathological verification, 78 pancreatic cancer specimens were subdivided into two groups: stage I (group 1) and stages II, III, and IV (group 2). We observed that the H-scores of COL1A2, HDGF, and CEBPD in stroma showed no significant variation in these two groups (**Figure 5F to 5H**). These results agree with the suggestion of HDGF and CEBPD in stroma contribute to pancreatic cancer associated-fibrosis.

**Figure 5.**
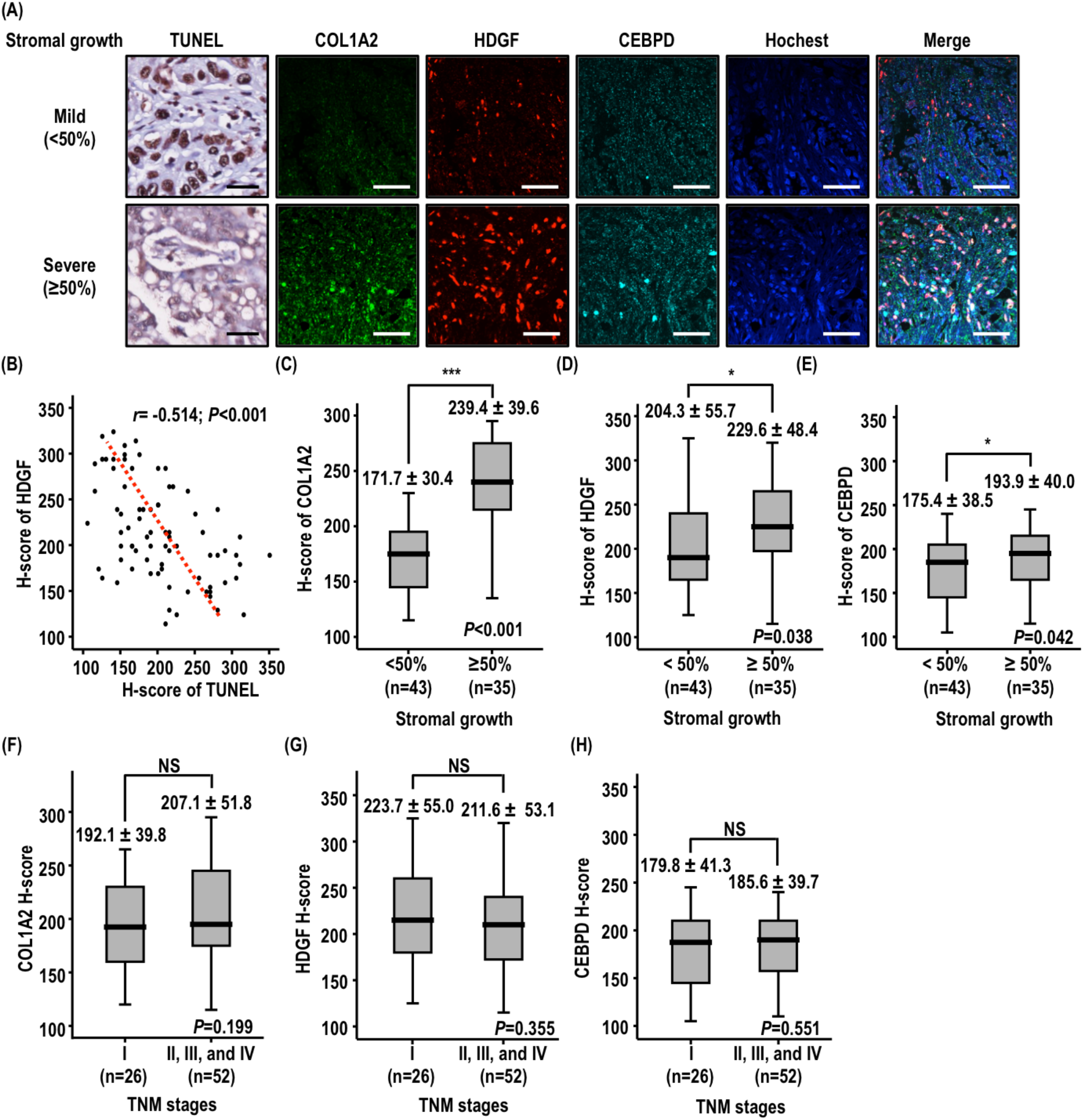
HDGF and CEBPD positively associate with anti-apoptosis and severe stromal growth in human pancreatic adenocarcinoma specimens. (A) The Opal^™^ multiplex tissue staining kit was used for immunofluorescence analysis on a commercial tissue array with 78 human primary pancreatic adenocarcinoma specimens. A ratio of cell proportions from stromal cells versus total tissue cells was determined. The 78 human primary pancreatic adenocarcinoma specimens were subdivided into mild (<50%) or severe (≥50%) stromal growth. Apoptotic cells were detected and labeled by brown color using TUNEL assay kit. Meanwhile, expression of COL1A2, HDGF, and CEBPD was determined in human pancreatic cancer specimens using immunofluorescence staining and labeled by green, red, and cyan fluorescence, respectively. The black and white scale bar indicated 100 μm. (B) to (E) Histochemistry-score (H-score), which assigned a score of 0-300 to each patient by an expert pathologist (Dr. Chien-Feng Li), is applicable to assess the extent of immunoreactivity of TUNEL (B), COL1A2 (C), HDGF (D), and CEBPD (E) in stroma. (B) Using Pearson’s correlation coefficient test, the negative and significant correlations were observed between stromal HDGF and apoptotic activity, which indicated cell apoptosis (*r*=-0.541; *P*<0.001). (C) to (E) The expression of COL1A2, HDGF, and CEBPD in stroma was significantly and positively associated with stromal overgrowth. The results are presented as both the mean ± standard error of mean and median with quartiles. Mann-Whitney U test was used for comparison between two groups, with P < 0.05 taken as indicating a significant difference (^*^ *P* < 0.05; ^**^ *P* < 0.01; ^***^ *P* < 0 .001). (F) to (H) According to Tumor/Node/Metastasis (TNM) staging, 78 pancreatic cancer specimens were grouped into two groups: stage I (group 1) and stages II, III, and IV (group 2). The H-scores of stromal COL1A2, HDGF, and CEBPD were comparable between two groups. The results are presented as both of mean ± standard error of mean and median with quartiles. Mann-Whitney U test was used for comparison between two groups, with P < 0.05 taken as indicating a significant difference (NS stands for not significant).

Our results suggested that TGF-β1-activated CEBPD/HIF-1α/HDFG axis in PSCs to involve in anti-apoptosis and pro-fibrosis under the normoxic conditions *in vitro*. However, most of clinical pancreatic cancer tissues are accompanied with strong fibrotic reactions, which associated with hypoxic status (Erkan et al., 2009). Therefore, to specifically assess whether the CEBPD/HIF-1α/HDFG axis promotes fibrosis under the hypoxic conditions, a hypoxic marker carbonic anhydrase IX (CA9) (Barathova, Takacova et al., 2008) was applied to distinguish the hypoxia/normoxia status of clinical specimens. Following the combination of CA9 abundance and the status of stromal growth, 78 pancreatic cancer specimens were subdivided into four groups as normoxic (low CA9 expression) or hypoxic (high CA9 expression) specimens of pancreatic adenocarcinoma accompanied with mild (<50%) or severe (≥50%) stromal growth (**Figure 6A and 6B**). The H-score of stromal CEBPD and HDGF were comparable between normoxic and hypoxic specimens of pancreatic adenocarcinoma (**Figure 6C and 6D**).We found that the higher H-score of stromal CEBPD and HDGF were observed and significantly associated with severe stromal growth in hypoxic specimens of pancreatic adenocarcinoma (**Figure 6E and 6F; compared groups 2 with groups 4**). Importantly, even in normoxic specimens of pancreatic adenocarcinoma, both the H-score of stromal CEBPD (from 181.4 ± 39.0 to 196.0 ±29.0) and stromal HDGF (from 211.6 ± 54.6 to 248.5 ± 45.1) were consistently elevated from mild to severe stromal growth (**Figure 6E and 6F; compared groups 1 with groups 3**). However, matter in normoxic or hypoxic specimens of pancreatic adenocarcinoma that accompanied severe stromal growth, there was no significant difference of the H-score of CEBPD and HDGF in stroma (**Figure 6E and 6F; compared groups 3 with groups 4**). Collectively, the above observations suggested that TGF-β1-activated CEBPD/HIF-1α/HDFG axis specifically associates with pancreatic cancer-associated fibrosis, but not is nothing to do with normoxia and hypoxia.

**Figure 6.**
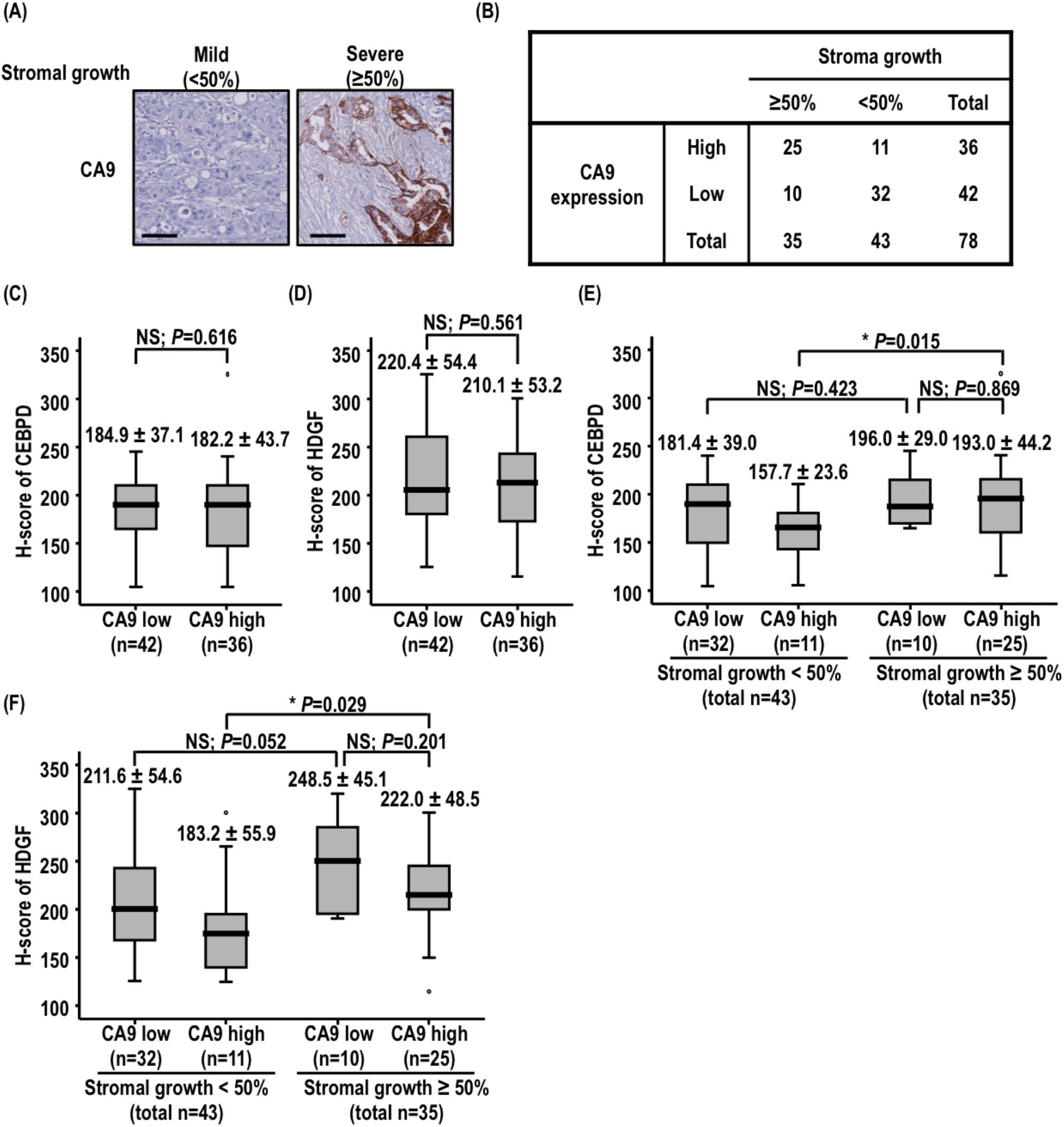
HDGF and CEBPD positively associate with anti-apoptosis and pro-fibrosis in normoxic and hypoxic human pancreatic adenocarcinoma specimens. (A) The antibody of carbonic anhydrase IX (CA9; as a hypoxic marker) was used for immunohistochemical analysis on a commercial tissue array with 78 human primary pancreatic adenocarcinoma specimens. A ratio of cell proportions from stromal cells versus total tissue cells was determined. The expression of CA9 (brown color) was assigned a H-score by an expert pathologist (CFL). The black scale bar indicated 200 μm. (B) Furthermore, according to the expression of CA9 and the status of stromal growth, 78 pancreatic cancer specimens were subdivided into four groups: normoxic (CA9 low expression) or hypoxic (CA9 high expression) specimens accompanied with mild (<50%) or severe (≥50%) stromal growth. (C) and (D) The expression of stromal CEBPD and HDGF showed comparable levels associated with stromal overgrowth in normoxic and hypoxic specimens. (E) and (F) The H-scores of stromal CEBPD and HDGF were selected and compared between two conditions among four sub-groups (normoxic or hypoxic specimens accompanied with mild or severe stromal growth). Above results are presented as both the mean ± standard error of mean and median with quartiles. Mann-Whitney U test was used for comparison between two groups, with P < 0.05 taken as indicating a significant difference (NS stands for not significant; circle dot indicated the mild outlier of H-score in the 78 human primary pancreatic adenocarcinoma specimens).

## DISCUSSION

The aim of the present study was to determine HDGF regulation and function in PSCs as well as to determine its contribution to fibrosis-associated pancreatic cancer. The results demonstrated that TGF-β1 activated HDGF expression through CEBPD/HIF-1α axis in PSCs. Increased production of HDGF prevented apoptosis and promoted the production of ECM proteins, including COL1A2 and FN1, to stabilize PSCs/PCCs tumor foci. Importantly, these results suggested that HDGF supported PSCs in orchestrating pancreatic cancer-associated fibrotic reactions, which act as a fibrotic hypo-vascular barrier to attenuate blood vessel penetration. In contrast, the attenuation of HDGF in PSCs resulted in less or looser fibrotic reactions, which could benefit outgrowth of PCCs (**Figure 7**).

**Figure 7.**
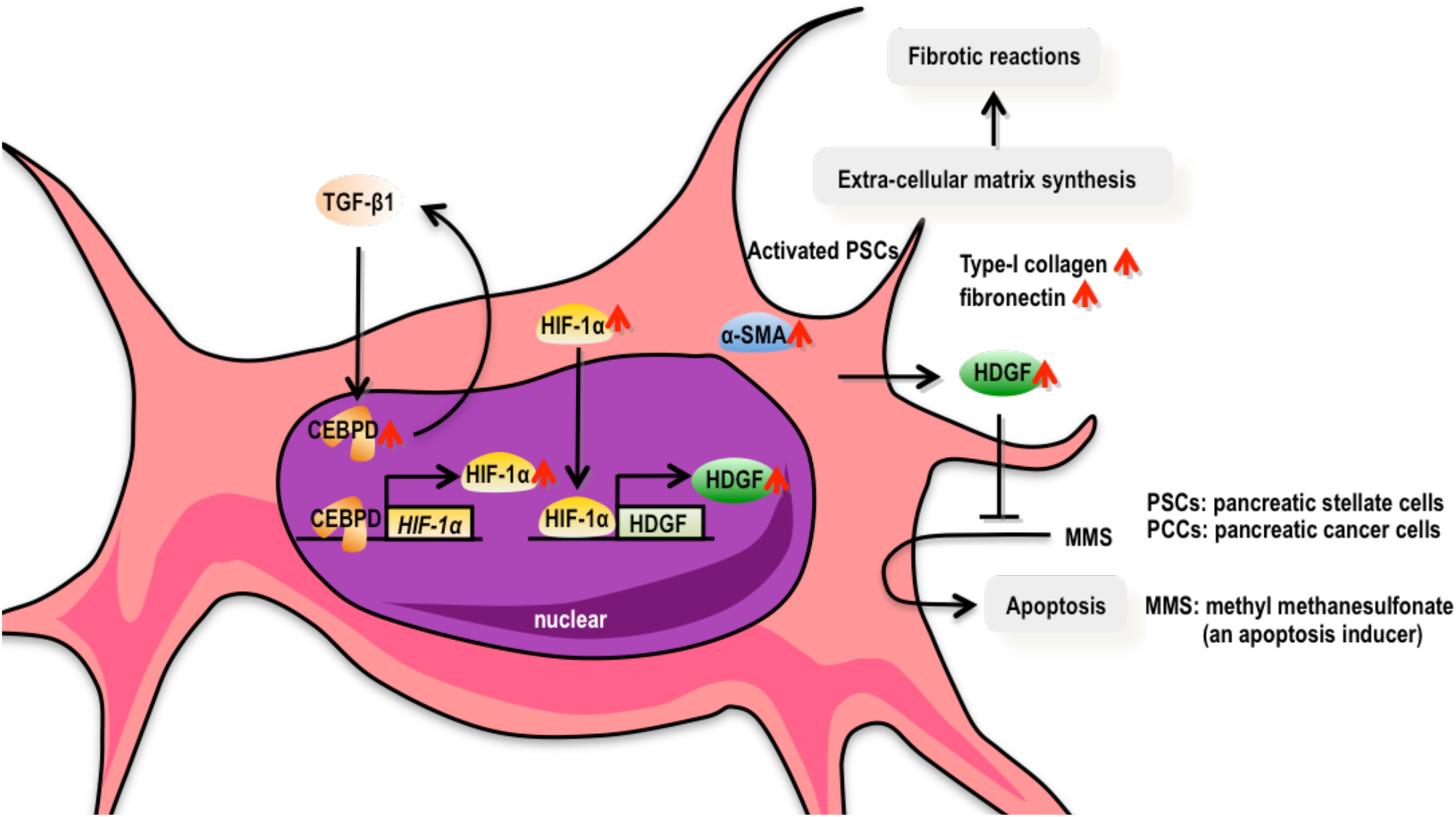
A schematic illustration of a signal pathway in pancreatic stellate cells to construct fibrotic reactions. TGF-β1-treated PSCs secreted HDGF through the TGF-β1/CEBPD/HIF-1α axis. The effects of HDGF in PSCs served anti-apoptosis under methyl methanesulfonate stimulation. Additionally, HDGF acted as a pro-fibrotic factor since its expression was required for TGF-β1-treated PSCs to synthesize and deposit extracellular matrix proteins, such as type-I collagen and fibronectin. These results suggest roles for HDGF in anti-apoptosis and pro-fibrosis in PSCs. Furthermore, TGF-β1 was induced by CEBPD through a reciprocal regulation. These results suggest that CEBPD serves as an upstream mediator to regulate HDGF expression in PSCs. Collectively, HDGF or CEBPD could be a therapeutic target to nullify the pancreatic cancer associated-fibrosis.

The evidence of HDGF auto-regulation and contribution to cell proliferation and tumorigenesis has been suggested in several types of cancer cells (Kishima et al., 2002, Song, Cong et al., 2017, Yang, Li et al., 2013). In the present study, we revealed that the regulation of HDGF in PSCs (**Figure 3**) contributed to anti-apoptosis and pro-fibrosis (**Figure 1 and 2**). The attenuation of HDGF in PSCs resulted in smaller and separated sphere formation of PCCs *in vitro* (**Figure 1H and 1I; Supplementary Figure 3C and 3D**). Moreover, the promoted tumor outgrowth of co-implanted *shHDGF #2* RLT-PSCs and EGFP-beating PCCs was observed in *in vivo* animal study (**Figure 2I, 2J, and Supplementary Figure 4**). These results in part agree with the observation that partially regaining physical restraint under less fibrotic reactions and blood irrigation to tumors through penetrating blood vessels (Rhim et al., 2014). Therefore, in addition to clarifying the individual contribution of HDGF from cancer cells and cancer-associated stellate cells, the most important issue should be to verify how to use our understanding of HDGF for pancreatic cancer diagnosis and therapy.

PSCs not only synthesize ECM proteins (Apte et al., 2012) but also provide pro-survival signals to tumors (Habisch et al., 2010, Masamune & Shimosegawa, 2009). Our results showed that HDGF knockdown RLT-PSCs orchestrated less or looser the fibrotic reactions in the co-implantation of PCCs and PSCs xenograft model (**Figure 2**). Interestingly, the signals of COL1A2 and FN1 were also observed in GFP-positive EGFP/PANC-1 or EGFP/MIA PaCa-2 cells in tissue sections (**Figure 2C to 2F**). Previous studies have discovered that pancreatic tumor cells express COL1A2 and FN1 (Lohr, Trautmann et al., 1994) and are able to take up collagen fragments from stromal microenvironment under nutrient limited conditions (Olivares, Mayers et al., 2017). We proposed that the accumulation of COL1A2 and FN1 in GFP-positive EGFP/PANC-1 or EGFP/MIA PaCa-2 cells could be resulted from above machinery. However, how stromal HDGF affects the cancerous expression of ECM proteins remains unrevealed.

Remarkably, fibrosis occurred in each TNM stage of pancreatic cancer (**Figure 5E**); no positive correlation was observed between the stromal HDGF expression or stromal CEBPD expression and malignancies of pancreatic cancers at various TNM stages (**Figure 5F and 5G**), suggesting that the fibrotic involvement of HDGF or CEBPD in the development of pancreatic cancer starts from an early stage with normoxia. A previous study suggests that precise quantitation of the fibrosis levels in pancreatic cancer should be combined to determine the stage of the disease, select optimal therapies, and monitor or predict patient therapeutic outcomes (Li, Zhang et al., 2013). The results of the present study suggest that HDGF and CEBPD contribute to the interplay of PSCs and PCCs in pancreatic cancer progression via depositions of ECM proteins, thereby indicating that HDGF or CEBPD can serve as a biomarker to evaluate the fibrotic status of pancreatic cancer.

Targeting tumors is currently a major tumor therapeutic strategy. Accumulating studies have highlighted the importance of the tumor microenvironment in benefiting cancer progression (Neesse, Algul et al., 2015, Tod, Jenei et al., 2013, Wilson, Pirola et al., 2014). Recent studies report that HGF secreted from pancreatic stromal cells controls pancreatic tumor metastasis through c-Met regulating tyrosine phosphorylation of Annexin A2 in PCCs (Rucki, Foley et al., 2017); concurrently, PCCs secrete Sonic Hedgehog to accelerate the growth of fibroblasts (Katagiri, Kobayashi et al., 2018). Disrupting the tumor-stroma interaction could be one of effective treatments for pancreatic cancer therapy. The results of the present study demonstrate that HDGF is involved in anti-apoptosis of PSCs in pancreatic cancer-associated fibrosis via the CEBPD/HIF-1α axis. However, two previous studies demonstrated that depletion of α-SMA-positive fibroblasts (Ozdemir et al., 2014) and loss of sonic hedgehog protein (Rhim et al., 2014) reduce fibrotic reactions and accelerate tumor progression, including enhanced tumor growth and angiogenesis, in undifferentiated and invasive tumors with reduced animal survival. Therefore, in the future, application of therapeutic strategies targeting CEBPD or HDGF in PSCs for reducing fibrotic reactions should be combined with chemotherapeutic drugs to kill pancreatic cancer cells.

## MATERIALS AND METHODS

### Cell culture

Immortalized human pancreatic stellate cells, RTL-PSCs, were generously provided from Dr. Kelvin K.-C. Tsai (National Health Research Institutes, Taiwan (Wang, Hsu et al., 2013)). RLT-PSCs were maintained in Dulbecco’s Modified Eagle’s medium (Life Technologies Co. Ltd., Grand Island, New York, United States; Gibco, #12900-082). Human pancreatic cancer cells, PANC-1 and MIA PaCa-2 cells, were generously provided from Dr. Yan-Shen Shan (National Cheng Kung University Hospital, Taiwan (Yang, Wang et al., 2015)). Both PANC-1 and MIA PaCa-2 cells were maintained in RPMI-1640 medium (GE Healthcare Life Sciences, Logan, Utah, United States; Hyclone, #SH30011.02). Phoenix™ ampho cells (purchased from Allele Biotechnology, San Diego, California, United States; #ABP-RVC-10001) for lenti-virus production were also maintained in Dulbecco’s Modified Eagle’s medium. All culture media contained 10% fetal bovine serum (Life Technologies Co. Ltd., Grand Island, New York, United States; Gibco, #10437-028) and penicillin-streptomycin solution (Mediatech, Inc., Manassas, Virginia, United States; Corning, #30-002-CI).

### Recombinant human TGF-β1, HDGF and chemical inhibitor reagents

Recombinant human TGF-β1 was purchased from the PeproTech, Inc. (Rocky Hill, New Jersey, United States; #100-21C). Recombinant human HDGF was kindly provided from Professor Ming-Hong Tai (National Sun Yat-sen University, Taiwan (Hu, Huang et al., 2003)). The protein synthesis inhibitor (cycloheximide, #C7698) was purchased from Sigma-Aldrich Co. (Saint Louis, Missouri, United States).

### Reverse transcription polymerase chain reaction (RT-PCR) assay

Total RNA was extracted from cells using TRIsure™ reagent (Bioline GmbH, Luckenwalde, Germany; #BIO-38033), and the RNA concentration was measured based on the absorbance at 260 nm using a Colibri Microvolume Spectrometer from Titertek-Berthold (Berthold Detection Systems GmbH, Pforzheim, Germany). cDNA was generated from 1 μg of RNA using SuperScript^®^ III Reverse Transcriptase according to the manufacturer’s instructions (Invitrogen Co., Carlsbad, California, United States; #18080085). The specific primer sequences of target gene amplification and reaction conditions are listed at **Supplementary Table 1**. For confirmation, the PCR product was electrophoresed on an agarose gel containing 100 mg/mL ethidium bromide.

### Protein extraction and western blot assays

Total proteins were extracted from cells using lysis buffer (10 mM of Tris-HC1 with pH 7.5, 150 mM of NaCl, 5 mM of EDTA, 5 mM of NaN_3_, 10 mM of sodium pyrophosphate, and 1% of Triton X-100). Subsequently, total proteins were separated using 8~15% polyacrylamide-sodium dodecyl sulfate gel. Total proteins were transferred onto an Immobilon^®^-P Transfer Membrane (Merck KGaA, Darmstadt, Germany; Millipore, #IPVH00010). Subsequently, the membrane was hybridized with a primary antibody (**Supplementary Table 2**) at 4°C overnight and a suitable secondary antibody (**Supplementary Table 2**) at room temperature for at least one hour. Finally, the UVP BioSpectrum 815 imaging system (UVP, LLC., Upland, California, United States) was used to detect the immunoreactivity of immune complexes after the membrane was incubated with Immobilon™ Western Chemiluminescent horseradish peroxidase substrate (Merck KGaA, Darmstadt, Germany; Millipore, #WBKLS0500).

### MTT assay for cell viability analysis

1×10^4^ of RLT-PSCs were seeded into 24-well plate and treated with or without TGF-β1 (0.5 ng/mL) for 24 hours, and then MTT assay was performed to measure TGF-β1 effects on cell proliferation. 200 μL of thiazolyl blue tetrazolium bromide, MTT (Sigma-Aldrich Co., Saint Louis, Missouri, United States; #M5655), with 0.5 mg/mL was added into each well and incubated for 4 hours. At the end of incubation, 500 μL of dimethyl sulfoxide per well was added to dissolve purple formazen and 200 μL of per well solution transferred into 96-well plate (Twentyman & Luscombe, 1987). The absorbance value at OD 595 nm (A595) was measured. The cell viability was calculated using the following formula: % cell viability = A595 of cells with or without TGF-β1/A595 of control cells without TGF-β1. The cell viability of RLT-PSCs without TGF-β1 was normalized to 100% as the control group.

### Propidium iodide (PI) staining for apoptosis analysis by flow cytometry

Methyl methanesulfonate (MMS; Sigma-Aldrich Co., Saint Louis, Missouri, United States; #129925) was used to induce cell death through apoptosis. Treated cells were fixed with 70% ethanol overnight and stained with propidium iodide (Sigma-Aldrich Co., Saint Louis, Missouri, United States; #P4170) to stain nuclear DNA at room temperature in the dark (thirty minutes). Using flow cytometry, a Cell Lab Quanta SC (Beckman Coulter, Inc., Brea, California, United States), 10,000 stained cells were acquired. The sub-G1 phase distribution, which indicated the apoptotic cell population, was analyzed on stained cells.

### Caspase-3/7activity for apoptosis analysis

After the medium of treated cells was removed, CellEvent™ caspase-3/7 green detection reagent (Thermo Fisher Scientific, Inc., Carlsbad, California, United States; Invitrogen, #C10423) was diluted into complete medium to a final concentration of 2 μM and incubated with treated cells at 37°C in dark for thirty minutes (final sample volume: 100 uL). Subsequently, the fluorogenic response was detected using Labsystems Finstruments 374 Fluoroskan Ascent Fluorescence microplate fluorometer and luminometer (Finland) with an excitation/emission of 485/538 nm.

### Lentiviral short hairpin (shRNA) system

Expression vectors (pLKO AS3w.puro and pLKO AS3w.eGFP.puro) and other target genes of shRNAs were purchased from the National RNAi Core Facility Platform of Academia Sinica in Taiwan, and the sequences are listed in **Supplementary Table 3**. Additionally, the *mCherry* fluorescent gene was inserted into the *Nhe*1 and *Pme*1 sites of the pLKO AS3w.puro vector. For lentivirus production, the expression vectors or shRNAs were transfected into Phoenix^™^ ampho cells using TransIT^®^-2020 Transfection reagent (Mirus Bio LLC., Madison, Wisconsin, United States) according to the manufacturer’s instructions. The lentivirus was purified, and the cells were transfected with the target genes of siRNAs using a lentivirus-based system according to the protocol on the website of the National RNAi Core Facility Platform of Academia Sinica. After lenti-viral infection, the cell lysate was extracted for western blot analysis or cells were subjected to fluorescence microscopy to determine the protein expression of the target gene. Subsequently, stable cell lines containing over-expressed or knocked-down target genes were established through antibiotics selection for subsequent experiments. In addition, cells with *shLacZ* (as a control group) or *shHDGF* were further transfected with the *mCherry* fluorescence gene. The method of limiting cell dilution was used to acquire monoclonal cells with the *mCherry* fluorescence gene.

### Hanging drop cell culture for three-dimension co-culture system of PSCs and PCCs

PANC-1 cells and MIA PaCa-2 cells were forced to express enhanced green fluorescence protein (EGFP). Subsequently, 2 × 10^4^ EGFP-bearing PANC-1 (EGFP/PANC-1) or EGFP-bearing MIA PaCa-2 (EGFP/MIA PaCa-2) cells were individually mixed with 1 × 10^5^ mCherry-bearing *shLacZ* RLT-PSCs (mCherry/*shLacZ* RLT-PSCs) or mCherry-bearing *shHDGF* RLT-PSCs (mCherry/*shHDGF* RLT-PSCs). The final volume of mixed cells was 400 μL, containing 50% matrigel matrix (Discovery Labware, Inc., Bedford, Massachusetts, United States; Corning, #356227). Several drops of the mixed cells were seeded onto 35 mm microplate with a glass bottom (ibidi GmbH, Martinsried, Germany; #81158) using hanging drop cell culture to generate a three-dimensional co-culture system. After 8 days, the mixed cells were observed using an Olympus FV1000 confocal microscope (Olympus Co., Shinjuku-ku, Tokyo, Japan) at 100 × magnification. Finally, the fluorescence intensity of the captured images was used to construct the three-dimensional structure according to the z-axis of the captured images using ImageJ software (National Institutes of Health, United States).

### Enzyme-linked immunosorbent (ELISA) assay

Human HDGF ELISA kit (Arigo Biolaboratories Co., Taiwan; #ARG81356) was applied to determine the levels of HDGF secretion according to the manual of the kit.. Briefly, 7×10^5^ of RLT-PSCs were seeded into 10 cm-dish. 5 mL of serum-free medium containing 1% bovine serum albumin with or without TGF-β1 (0.5 ng/mL) was used for 24 hours of experimental treatment. The tested samples were made after cell medium collection, centrifugation at 1,500 rpm. After incubating for 2 hours, tested samples were added to 100 μL HRP-antibody conjugate and incubated for 1 hour in dark, subsequently washed with wash buffer, and incubated in substrate solution for 10 minutes in dark. Finally, optical density at 450 nm was read after 50 μL of stop solution was added and gently mixed. A standard curve was generated from antigen standard wells. Duplicate wells were run for all standards and tested samples. The results of the mean absorbance value were calculated automatically using 4 parameter logistics curve fit.

### Transcription factor prediction

The 2000 bases of the promoter sequences (upstream bases from -2000 to -1 before transcript start site) of HIF-1α and CEBPD were acquired from the University of California Santa Cruz Genome Browser. The promoter sequence was analyzed using the TFSEARCH (version 1.3) and PROMO server (version 3.0.2), for transcription factor prediction. All the University of California Santa Cruz Genome Browser, TFSEARCH (version 1.3), and PROMO server (version 3.0.2) are open database sources. The predicted transcription factors were further verified using luciferase reporter and chromatin immunoprecipitation (ChIP) assays.

### Plasmid transfection

Human HIF-1α expressing vector (pCEP4/HIF-1α; American type Culture Collection, Manassas, Virginia, United States; #MBA-2) was a generous gift from Professor Shaw-Jenq Tsai (National Cheng Kung University (NCKU), Taiwan) (Chien, Lin et al., 2008). Furthermore, the pcDNA3/HA-CEBPD expressing vector was established previously described (Lai, Wang et al., 2008). Plasmids were transfected into cells using the Turbofect^™^ Transfection reagent (Thermo Fisher Scientific, Inc., Rockford, Illinois, United States; #R0532). Subsequently, the cell lysate was extracted and subjected to western blot analysis to determine the protein expression of the target gene.

### Reporter plasmid construction and luciferase reporter assay

Two promoter fragments of the human *HDGF* gene cloned into pGL3-Enhancer vectors (pHDGF 2047: -2017 to +30 and pHDGF 876: -774 to +102) were kindly provided from Professor Ming-Hong Tai (National Sun Yat-sen University, Taiwan). In addition, we also designed appropriate primers to generate two promoter fragments of the human *HIF-1α* gene; the primer sequences are listed in **Supplementary Table 4**. Briefly, genomic DNA was extracted from cells using the GeneJET^™^ Genomic DNA Purification Kit (Thermo Fisher Scientific, Inc., Waltham, Massachusetts, United States; Fermentas, #K0722) and used as a template to generate promoter fragments of the target genes using PCR. The promoter fragments of the *HIF-1α* gene were subsequently cloned into pGL3-Basic vectors (Promega Co., Madison, Wisconsin, United States; #E1751). After verification by sequencing, the pGL3-Enhancer vectors bearing promoter fragments of the *HDGF* gene or pGL3-Basic vectors bearing promoter fragments of the *HIF-1α* gene were transfected into cells using the Turbofect™ Transfection reagent. Subsequently, the cell lysate was extracted and analyzed using a luciferase assay system.

### Chromatin immunoprecipitation (ChIP) assay

ChIP assay was used to orchestrate protein-DNA interactions. Briefly, the crosslink between DNA and associated proteins on chromatin in living cells was fixed using 1% formaldehyde and neutralized using 1.375 M Glycine. Subsequently, DNA-protein complexes (chromatin-protein) were broken, and DNA fragments shorter less than 1000 base pairs were generated using the Misonix Sonicator 3000 Ultrasonic Cell Disruptor (Misonix, Inc., Farmingdale, New York, United States). The cross-linked DNA fragments associated with protein were selectively immuno-precipitated using primary antibodies as listed in **Supplementary Table 2** and Protein A Agarose/Salmon Sperm DNA beads (Merck KGaA, Darmstadt, Germany; Upstate, #16-157). The associated DNA fragments were purified and subjected to PCR using the appropriate primers as listed in **Supplementary Table 5** for putative protein binding motifs. Finally, the PCR product was confirmed using agarose gel electrophoresis with ethidium bromide staining.

### Animal xenograft model for co-implantation of RLT-PSCs and PCCs

The animal experiments were approved by the Institutional Animal Care and Use Committee of NCKU (IACUC number 03239). Non-obese diabetic-severe combined immunodeficiency (NOD-SCID) male mice were acquired and maintained in the specific pathogen-free environment at the Laboratory Animal Center of College of Medicine of NCKU. *shLacZ* or *shHDGF* RLT-PSCs were individually mixed with EGFP/PANC-1 or EGFP/MIA PaCa-2 cells (each at 1 × 10^6^ cells/50 μL). The mixed cells were added to 100 μL of matrigel matrix (final concentration: 50%; total final volume: 200 μL) and subcutaneously injected into the right flank of 7-week old NOD-SCID mice. During the co-inoculation of RLT-PSCs and PCCs for nine weeks, *in vivo* imaging system spectrum was applied to evaluate subcutaneous tumor size. After mice being sacrificed, subcutaneous tumor tissues were dissected and sent to the Human Biobank Research Center of Clinical Medicine of NCKU Hospital for fixing, paraffin-embedding and sectioning. The 4-μm sections of paraffin-embedded tumor tissues were applied for the following experiments.

### Picrosirius red staining for collagen analysis

Picrosirius red stain is specific with respect different ECM proteins and materials, such as collagens, laminin, fibronectin, chondroitin sulfate, dermatan sulfate, and amyloid β (Tullberg-Reinert & Jundt, 1999). After the tissue sections were de-paraffinized, a Picrosirius red stain kit (Abcam plc., Cambridge, United Kingdom; #ab150681) was used to stain collagen expression in tumor sections according to the manufacturer’s instructions. Images of the collagen histology were visualized and captured using an OLYMPUS DP70 microscope.

### Immunohistochemistry (IHC) analysis

All tissue sections were de-paraffinized in 100% xylene and a gradient alcohol concentration. The Novolink^™^ Polymer Detection Systems Kit (Leica Biosystems Newcastle Ltd., Benton, Lane, Newcastle Upon Tyne, United Kingdom; #RE7140-K) was used to perform IHC analysis according to the manufacturer’s instructions. Antigen retrieval was performed in Tris-EDTA buffer at 95°C for fifteen minutes. Subsequently, the tissue sections were incubated with primary antibody at 4°C overnight. The dilutions of the primary antibodies depended on the antigen and are listed in **Supplementary Table 2**. The peroxidase activity was developed using a DAB working solution. Finally, hematoxylin solution (Sigma-Aldrich Co., Saint Louis, Missouri, United States; #HHS32) was used for counter-staining. All images of the tissue sections were observed and captured using an OLYMPUS DP70 microscope.

### Immunofluorescence (IF) analysis

Cells and 4-μm of the paraffin-embedded mouse sections were used for IF analysis. Cells were fixed with 4% paraformaldehyde. After the tissue sections were de-paraffinized, Tris-EDTA buffer (10 mM Tris, 1 mM EDTA, and 0.05% Tween 20, pH 9) was used to perform antigen retrieval at 95°C for fifteen minutes. The primary and secondary antibodies were diluted depending on the various antigens listed in **Supplementary Table 2**. Subsequently, fixed cells or de-paraffinized mouse tissue sections were incubated with primary antibody at 4°C overnight and a suitable fluorescence secondary antibody at room temperature in dark for 1 hour. Finally, Hoechst 33258 (Sigma-Aldrich Co., Saint Louis, Missouri, United States; #861405) was used for nuclear DNA staining. Phalloidin-TRITC labeled (Sigma-Aldrich Co., Saint Louis, Missouri, United States; #77418) was also applied for filamentous actin (F-actin) staining on fixed cells for 1 hour. The reagent of ProLong^™^ diamond antifade mountant with DAPI (Thermo Fisher Scientific, Inc., Waltham, Massachusetts, United States; Life technologies, #P36962) was used for nuclear DNA staining and cell mounting. Finally, images of the cells or tissue sections were observed and captured using an Olympus DP70 microscope or FV10i confocal microscope (Olympus Co., Shinjuku-ku, Tokyo, Japan).

Additionally, the Opal^™^ multiplex tissue staining kit (PerkinElmer, Inc., Boston, Massachusetts, United States; #NEL794001KT) was applied for IF analysis of a commercial tissue array with 78 human primary pancreatic adenocarcinoma specimens (purchased from US Biomax, Inc., Derwood, Maryland, United States; # PA961b). The primary and secondary antibodies were diluted depending on the antigens and are listed in **Supplementary Table 2**. The fluorescent intensity and positive staining cell proportion of total cells were evaluated by an expert pathologist (Dr. Chien-Feng Li) and presented as a corresponding histochemistry score (H-score) as previously described (Rezaeian, Li et al., 2017). Briefly, H-score method for assessing the extent of immunoreactivity assigns a score of 0-300 to each patient. Experiments performed on human samples were approved by the institutional review board of Chi Mei Medical Center (10606-E03).

### TdT-mediated dUTP nick end labeling (TUNEL) assay

The TUNEL assay was performed as previously described (Zhang, Li et al., 2016).The TUNEL kit was purchased from Roche Diagnostics Deutschland GmbH (Germany; #1168479510) and applied for detection of fragmented DNA in apoptotic cells. Briefly, following deparaffinization and proteinase K treatment of the tissue sections of subcutaneous tumors and human primary pancreatic adenocarcinoma specimens, the tissue sections were incubated with TUNEL reaction mixture at 37°C for 1 hour. TUNEL-positive cells were scored by an expert pathologist (Dr. Chien-Feng Li).

### Statistical analysis

GraphPad Prism 5 software (GraphPad Software, Inc., La Jolla, California, United States) was used to assess statistical significance using Student’s unpaired T test in cell-based *in vitro* assays . The statistical significance was assessed in the animal study *in vivo* using Student’s unpaired T test or Mann-Whitney U test. The clinical specimens were assessed statistical significance using Mann-Whitney U test or statistical correlation using Pearson’s correlation coefficient test. The data are presented as the means ± standard error of the mean or median with quartiles. *P* values less than 0.05 were considered statistically significant and are denoted by an asterisk, and NS stands for not significant (^*^: *P* < 0.05; ^**^: *P* < 0.01; ^***^: *P* < 0.001; ^****^: *P* < 0.0001).

## ACKNOWLEDGMENTS

This work was financially supported by the National Science Council Grant (NHRI-EX106-10422BI from the National Health Research Institute and MOST 106-2320-B-006-063-MY3 from the Ministry of Science and Technology) and Chi Mei Medical Center Research Grant #10501. We are grateful for the experimental material providing from Dr. Kelvin K.-C. Tsai (National Health Research Institutes, Taiwan), Dr. Yan-Shen Shan (National Cheng Kung University Hospital, Taiwan), Professor Shaw-Jenq Tsai (National Cheng Kung University, Taiwan), and Professor Ming-Hong Tai (National Sun Yat-sen University, Taiwan).

## AUTHOR CONTRIBUTIONS

Y.-T. Chen were responsible to design and perform most of the experiments. Meanwhile, Y.-T. Chen analyzed and interpreted the data and results for writing this manuscript. C.-F. Li performed the clinical human studies and provided helpful suggestions for designing the experiments and editing the manuscript. T.-W. Wang constructed the reporter plasmids, and performed luciferase reporter assays, and assisted with part of immunofluorescence staining assays. T.-H. Chang assisted with the animal studies and the manuscript editing. T.-P. Hsu performed the chromatin immunoprecipitation assays, and assisted with the lenti-virus production. J.-Y. Chi and Y.-W. Hsiao cloned *mCherry* fluorescent gene into the pLKO AS3w.puro expression vector. J.-Y. Chi and Y.-W. Hsiao also supervised and assisted with T.-W. Wang’s experiments. J.-M. Wang directed the project and edited the manuscript.

## REFERENCES

Apte MV, Pirola RC, Wilson JS (2012) Pancreatic stellate cells: a starring role in normal and diseased pancreas. Front Physiol 3: 344

Bachem MG, Schneider E, Gross H, Weidenbach H, Schmid RM, Menke A, Siech M, Beger H, Grunert A, Adler G (1998) Identification, culture, and characterization of pancreatic stellate cells in rats and humans. Gastroenterology 115: 421-32

Bao C, Wang J, Ma W, Wang X, Cheng Y (2014) HDGF: a novel jack-of-all-trades in cancer. Future Oncol 10: 2675-85

Barathova M, Takacova M, Holotnakova T, Gibadulinova A, Ohradanova A, Zatovicova M, Hulikova A, Kopacek J, Parkkila S, Supuran CT, Pastorekova S, Pastorek J (2008) Alternative splicing variant of the hypoxia marker carbonic anhydrase IX expressed independently of hypoxia and tumour phenotype. Br J Cancer 98: 129-36

Binenbaum Y, Na’ara S, Gil Z (2015) Gemcitabine resistance in pancreatic ductal adenocarcinoma. Drug resistance updates : reviews and commentaries in antimicrobial and anticancer chemotherapy 23: 55-68

Burris HA, 3rd, Moore MJ, Andersen J, Green MR, Rothenberg ML, Modiano MR, Cripps MC, Portenoy RK, Storniolo AM, Tarassoff P, Nelson R, Dorr FA, Stephens CD, Von Hoff DD (1997) Improvements in survival and clinical benefit with gemcitabine as first-line therapy for patients with advanced pancreas cancer: a randomized trial. Journal of clinical oncology : official journal of the American Society of Clinical Oncology 15: 2403-13

Chen SC, Kung ML, Hu TH, Chen HY, Wu JC, Kuo HM, Tsai HE, Lin YW, Wen ZH, Liu JK, Yeh MH, Tai MH (2012) Hepatoma-derived growth factor regulates breast cancer cell invasion by modulating epithelial--mesenchymal transition. J Pathol 228: 158-69

Chi JY, Hsiao YW, Li CF, Lo YC, Lin ZY, Hong JY, Liu YM, Han X, Wang SM, Chen BK, Tsai KK, Wang JM (2015) Targeting chemotherapy-induced PTX3 in tumor stroma to prevent the progression of drug-resistant cancers. Oncotarget 6: 23987-4001

Chien CW, Lin SC, Lai YY, Lin BW, Lin SC, Lee JC, Tsai SJ (2008) Regulation of CD151 by hypoxia controls cell adhesion and metastasis in colorectal cancer. Clinical cancer research : an official journal of the American Association for Cancer Research 14: 8043-51

Chuang CH, Wang WJ, Li CF, Ko CY, Chou YH, Chuu CP, Cheng TL, Wang JM (2014) The combination of the prodrugs perforin-CEBPD and perforin-granzyme B efficiently enhances the activation of caspase signaling and kills prostate cancer. Cell Death Dis 5: e1220

Conroy T, Desseigne F, Ychou M, Bouche O, Guimbaud R, Becouarn Y, Adenis A, Raoul JL, Gourgou-Bourgade S, de la Fouchardiere C, Bennouna J, Bachet JB, Khemissa-Akouz F, Pere-Verge D, Delbaldo C, Assenat E, Chauffert B, Michel P, Montoto-Grillot C, Ducreux M et al. (2011) FOLFIRINOX versus gemcitabine for metastatic pancreatic cancer. N Engl J Med 364: 1817-25

Enomoto H, Nakamura H, Liu W, Nishiguchi S (2015) Hepatoma-Derived Growth Factor: Its Possible Involvement in the Progression of Hepatocellular Carcinoma. Int J Mol Sci 16: 14086-97

Erkan M, Reiser-Erkan C, Michalski CW, Deucker S, Sauliunaite D, Streit S, Esposito I, Friess H, Kleeff J (2009) Cancer-stellate cell interactions perpetuate the hypoxia-fibrosis cycle in pancreatic ductal adenocarcinoma. Neoplasia 11: 497-508

Fu Y, Liu S, Zeng S, Shen H (2018) The critical roles of activated stellate cells-mediated paracrine signaling, metabolism and onco-immunology in pancreatic ductal adenocarcinoma. Mol Cancer 17: 62

Habisch H, Zhou S, Siech M, Bachem MG (2010) Interaction of stellate cells with pancreatic carcinoma cells. Cancers (Basel) 2: 1661-82

Hsiao YW, Li CF, Chi JY, Tseng JT, Chang Y, Hsu LJ, Lee CH, Chang TH, Wang SM, Wang DD, Cheng HC, Wang JM (2013) CCAAT/enhancer binding protein delta in macrophages contributes to immunosuppression and inhibits phagocytosis in nasopharyngeal carcinoma. Sci Signal 6: ra59

Hu TH, Huang CC, Liu LF, Lin PR, Liu SY, Chang HW, Changchien CS, Lee CM, Chuang JH, Tai MH (2003) Expression of hepatoma-derived growth factor in hepatocellular carcinoma. Cancer 98: 1444-56

Jacobetz MA, Chan DS, Neesse A, Bapiro TE, Cook N, Frese KK, Feig C, Nakagawa T, Caldwell ME, Zecchini HI, Lolkema MP, Jiang P, Kultti A, Thompson CB, Maneval DC, Jodrell DI, Frost GI, Shepard HM, Skepper JN, Tuveson DA (2013) Hyaluronan impairs vascular function and drug delivery in a mouse model of pancreatic cancer. Gut 62: 112-20

Jesnowski R, Furst D, Ringel J, Chen Y, Schrodel A, Kleeff J, Kolb A, Schareck WD, Lohr M (2005) Immortalization of pancreatic stellate cells as an in vitro model of pancreatic fibrosis: deactivation is induced by matrigel and N-acetylcysteine. Lab Invest 85: 1276-91

Kao YH, Chen CL, Jawan B, Chung YH, Sun CK, Kuo SM, Hu TH, Lin YC, Chan HH, Cheng KH, Wu DC, Goto S, Cheng YF, Chao D, Tai MH (2010) Upregulation of hepatoma-derived growth factor is involved in murine hepatic fibrogenesis. Journal of hepatology 52: 96-105

Katagiri T, Kobayashi M, Yoshimura M, Morinibu A, Itasaka S, Hiraoka M, Harada H (2018) HIF-1 maintains a functional relationship between pancreatic cancer cells and stromal fibroblasts by upregulating expression and secretion of Sonic hedgehog. Oncotarget 9: 10525-10535

Kishima Y, Yamamoto H, Izumoto Y, Yoshida K, Enomoto H, Yamamoto M, Kuroda T, Ito H, Yoshizaki K, Nakamura H (2002) Hepatoma-derived growth factor stimulates cell growth after translocation to the nucleus by nuclear localization signals. J Biol Chem 277: 10315-22

Ko CY, Chang WC, Wang JM (2015) Biological roles of CCAAT/Enhancer-binding protein delta during inflammation. J Biomed Sci 22: 6

Ko CY, Hsu HC, Shen MR, Chang WC, Wang JM (2008) Epigenetic silencing of CCAAT/enhancer-binding protein delta activity by YY1/polycomb group/DNA methyltransferase complex. J Biol Chem 283: 30919-32

Kota J, Hancock J, Kwon J, Korc M (2017) Pancreatic cancer: Stroma and its current and emerging targeted therapies. Cancer Lett 391: 38-49

Lai PH, Wang WL, Ko CY, Lee YC, Yang WM, Shen TW, Chang WC, Wang JM (2008) HDAC1/HDAC3 modulates PPARG2 transcription through the sumoylated CEBPD in hepatic lipogenesis. Biochimica et biophysica acta 1783: 1803-14

Li CF, Tsai HH, Ko CY, Pan YC, Yen CJ, Lai HY, Yuh CH, Wu WC, Wang JM (2015) HMDB and 5-AzadC Combination Reverses Tumor Suppressor CCAAT/Enhancer-Binding Protein Delta to Strengthen the Death of Liver Cancer Cells. Mol Cancer Ther 14: 2623-33

Li W, Zhang Z, Nicolai J, Yang GY, Omary RA, Larson AC (2013) Quantitative magnetization transfer MRI of desmoplasia in pancreatic ductal adenocarcinoma xenografts. NMR in biomedicine 26: 1688-95

Li X, Ma Q, Xu Q, Duan W, Lei J, Wu E (2012) Targeting the cancer-stroma interaction: a potential approach for pancreatic cancer treatment. Curr Pharm Des 18: 2404-15

Lohr M, Trautmann B, Gottler M, Peters S, Zauner I, Maillet B, Kloppel G (1994) Human ductal adenocarcinomas of the pancreas express extracellular matrix proteins. Br J Cancer 69: 144-51

Majmundar AJ, Wong WJ, Simon MC (2010) Hypoxia-inducible factors and the response to hypoxic stress. Mol Cell 40: 294-309

Masamune A, Kikuta K, Watanabe T, Satoh K, Hirota M, Shimosegawa T (2008) Hypoxia stimulates pancreatic stellate cells to induce fibrosis and angiogenesis in pancreatic cancer. Am J Physiol Gastrointest Liver Physiol 295: G709-17

Masamune A, Shimosegawa T (2009) Signal transduction in pancreatic stellate cells. J Gastroenterol 44: 249-60

Masamune A, Watanabe T, Kikuta K, Shimosegawa T (2009) Roles of pancreatic stellate cells in pancreatic inflammation and fibrosis. Clin Gastroenterol Hepatol 7: S48-54

McCarroll JA, Naim S, Sharbeen G, Russia N, Lee J, Kavallaris M, Goldstein D, Phillips PA (2014) Role of pancreatic stellate cells in chemoresistance in pancreatic cancer. Front Physiol 5: 141

Menke A, Adler G (2002) TGFbeta-induced fibrogenesis of the pancreas. Int J Gastrointest Cancer 31: 41-6

Michl P, Gress TM (2012) Improving drug delivery to pancreatic cancer: breaching the stromal fortress by targeting hyaluronic acid. Gut 61: 1377-9

Nakamura H, Izumoto Y, Kambe H, Kuroda T, Mori T, Kawamura K, Yamamoto H, Kishimoto T (1994) Molecular cloning of complementary DNA for a novel human hepatoma-derived growth factor. Its homology with high mobility group-1 protein. The Journal of biological chemistry 269: 25143-9

Neesse A, Algul H, Tuveson DA, Gress TM (2015) Stromal biology and therapy in pancreatic cancer: a changing paradigm. Gut 64: 1476-84

Olivares O, Mayers JR, Gouirand V, Torrence ME, Gicquel T, Borge L, Lac S, Roques J, Lavaut MN, Berthezene P, Rubis M, Secq V, Garcia S, Moutardier V, Lombardo D, Iovanna JL, Tomasini R, Guillaumond F, Vander Heiden MG, Vasseur S (2017) Collagen-derived proline promotes pancreatic ductal adenocarcinoma cell survival under nutrient limited conditions. Nat Commun 8: 16031

Olive KP, Jacobetz MA, Davidson CJ, Gopinathan A, McIntyre D, Honess D, Madhu B, Goldgraben MA, Caldwell ME, Allard D, Frese KK, Denicola G, Feig C, Combs C, Winter SP, Ireland-Zecchini H, Reichelt S, Howat WJ, Chang A, Dhara M et al. (2009) Inhibition of Hedgehog signaling enhances delivery of chemotherapy in a mouse model of pancreatic cancer. Science 324: 1457-61

Ooi BN, Mukhopadhyay A, Masilamani J, Do DV, Lim CP, Cao XM, Lim IJ, Mao L, Ren HN, Nakamura H, Phan TT (2010) Hepatoma-derived growth factor and its role in keloid pathogenesis. J Cell Mol Med 14: 1328-37

Ozdemir BC, Pentcheva-Hoang T, Carstens JL, Zheng X, Wu CC, Simpson TR, Laklai H, Sugimoto H, Kahlert C, Novitskiy SV, De Jesus-Acosta A, Sharma P, Heidari P, Mahmood U, Chin L, Moses HL, Weaver VM, Maitra A, Allison JP, LeBleu VS et al. (2014) Depletion of carcinoma-associated fibroblasts and fibrosis induces immunosuppression and accelerates pancreas cancer with reduced survival. Cancer cell 25: 719-34

Rezaeian AH, Li CF, Wu CY, Zhang X, Delacerda J, You MJ, Han F, Cai Z, Jeong YS, Jin G, Phan L, Chou PC, Lee MH, Hung MC, Sarbassov D, Lin HK (2017) A hypoxia-responsive TRAF6-ATM-H2AX signalling axis promotes HIF1alpha activation, tumorigenesis and metastasis. Nat Cell Biol 19: 38-51

Rhim AD, Oberstein PE, Thomas DH, Mirek ET, Palermo CF, Sastra SA, Dekleva EN, Saunders T, Becerra CP, Tattersall IW, Westphalen CB, Kitajewski J, Fernandez-Barrena MG, Fernandez-Zapico ME, Iacobuzio-Donahue C, Olive KP, Stanger BZ (2014) Stromal elements act to restrain, rather than support, pancreatic ductal adenocarcinoma. Cancer Cell 25: 735-47

Rucki AA, Foley K, Zhang P, Xiao Q, Kleponis J, Wu AA, Sharma R, Mo G, Liu A, Van Eyk J, Jaffee EM, Zheng L (2017) Heterogeneous Stromal Signaling within the Tumor Microenvironment Controls the Metastasis of Pancreatic Cancer. Cancer Res 77: 41-52

Sakurai Y, Sawada T, Chung YS, Funae Y, Sowa M (1997) Identification and characterization of motility stimulating factor secreted from pancreatic cancer cells: role in tumor invasion and metastasis. Clin Exp Metastasis 15: 307-17

Shields MA, Dangi-Garimella S, Redig AJ, Munshi HG (2012) Biochemical role of the collagen-rich tumour microenvironment in pancreatic cancer progression. Biochem J 441: 541-52

Shih TC, Tien YJ, Wen CJ, Yeh TS, Yu MC, Huang CH, Lee YS, Yen TC, Hsieh SY (2012) MicroRNA-214 downregulation contributes to tumor angiogenesis by inducing secretion of the hepatoma-derived growth factor in human hepatoma. J Hepatol 57: 584-91

Song R, Cong L, Ni G, Chen M, Sun H, Sun Y, Chen M (2017) MicroRNA-195 inhibits the behavior of cervical cancer tumors by directly targeting HDGF. Oncol Lett 14: 767-775

Suetsugu A, Snyder CS, Moriwaki H, Saji S, Bouvet M, Hoffman RM (2015) Imaging the Interaction of Pancreatic Cancer and Stellate Cells in the Tumor Microenvironment during Metastasis. Anticancer Res 35: 2545-51

Tod J, Jenei V, Thomas G, Fine D (2013) Tumor-stromal interactions in pancreatic cancer. Pancreatology 13: 1-7

Tullberg-Reinert H, Jundt G (1999) In situ measurement of collagen synthesis by human bone cells with a sirius red-based colorimetric microassay: effects of transforming growth factor beta2 and ascorbic acid 2-phosphate. Histochemistry and cell biology 112: 271-6

Twentyman PR, Luscombe M (1987) A study of some variables in a tetrazolium dye (MTT) based assay for cell growth and chemosensitivity. Br J Cancer 56: 279-85

Vogelmann R, Ruf D, Wagner M, Adler G, Menke A (2001) Effects of fibrogenic mediators on the development of pancreatic fibrosis in a TGF-beta1 transgenic mouse model. Am J Physiol Gastrointest Liver Physiol 280: G164-72

Von Hoff DD, Ervin T, Arena FP, Chiorean EG, Infante J, Moore M, Seay T, Tjulandin SA, Ma WW, Saleh MN, Harris M, Reni M, Dowden S, Laheru D, Bahary N, Ramanathan RK, Tabernero J, Hidalgo M, Goldstein D, Van Cutsem E et al. (2013) Increased survival in pancreatic cancer with nab-paclitaxel plus gemcitabine. N Engl J Med 369: 1691-703

Vonlaufen A, Phillips PA, Yang L, Xu Z, Fiala-Beer E, Zhang X, Pirola RC, Wilson JS, Apte MV (2010) Isolation of quiescent human pancreatic stellate cells: a promising in vitro tool for studies of human pancreatic stellate cell biology. Pancreatology 10: 434-43

Wang WY, Hsu CC, Wang TY, Li CR, Hou YC, Chu JM, Lee CT, Liu MS, Su JJ, Jian KY, Huang SS, Jiang SS, Shan YS, Lin PW, Shen YY, Lee MT, Chan TS, Chang CC, Chen CH, Chang IS et al. (2013) A gene expression signature of epithelial tubulogenesis and a role for ASPM in pancreatic tumor progression. Gastroenterology 145: 1110-20

Wehr AY, Furth EE, Sangar V, Blair IA, Yu KH (2011) Analysis of the human pancreatic stellate cell secreted proteome. Pancreas 40: 557-66

Wilson JS, Pirola RC, Apte MV (2014) Stars and stripes in pancreatic cancer: role of stellate cells and stroma in cancer progression. Front Physiol 5: 52

Yamamoto S, Tomita Y, Hoshida Y, Takiguchi S, Fujiwara Y, Yasuda T, Doki Y, Yoshida K, Aozasa K, Nakamura H, Monden M (2006) Expression of hepatoma-derived growth factor is correlated with lymph node metastasis and prognosis of gastric carcinoma. Clin Cancer Res 12: 117-22

Yang MC, Wang HC, Hou YC, Tung HL, Chiu TJ, Shan YS (2015) Blockade of autophagy reduces pancreatic cancer stem cell activity and potentiates the tumoricidal effect of gemcitabine. Mol Cancer 14: 179

Yang Y, Li H, Zhang F, Shi H, Zhen T, Dai S, Kang L, Liang Y, Wang J, Han A (2013) Clinical and biological significance of hepatoma-derived growth factor in Ewing’s sarcoma. J Pathol 231: 323-34

Zhang X, Li CF, Zhang L, Wu CY, Han L, Jin G, Rezaeian AH, Han F, Liu C, Xu C, Xu X, Huang CY, Tsai FJ, Tsai CH, Watabe K, Lin HK (2016) TRAF6 Restricts p53 Mitochondrial Translocation, Apoptosis, and Tumor Suppression. Mol Cell 64: 803-814

